# A Mem-dELISA platform for dual color and ultrasensitive digital detection of colocalized proteins on extracellular vesicles

**DOI:** 10.1101/2024.03.10.584311

**Authors:** Himani Sharma, Vivek Yadav, Alice Burchett, Tiger Shi, Satyajyoti Senapati, Meenal Datta, Hsueh-Chia Chang

## Abstract

Accurate, multiplex, and ultrasensitive measurement of different colocalized protein markers on individual tumor-derived extracellular vesicles (EVs) and dimerized proteins with multiple epitopes could provide insights into cancer heterogeneity, therapy management and early diagnostics that cannot be extracted from bulk methods. However, current digital protein assays lack certain features to enable robust colocalization, including multi-color detection capability, large dynamic range, and selectivity against background proteins. Here, we report a lithography-free, inexpensive (< $0.1) and ultrasensitive dual-color Membrane Digital ELISA (Mem-dELISA) platform by using track-etched polycarbonate (PCTE) membranes to overcome these shortcomings. Their through-pores remove air bubbles through wicking before they are sealed on one side by adhesion to form microwells. Immunomagnetic bead-analyte complexes and substrate solution are then loaded into the microwells from the opposite side, with >80% loading efficiency, before sealing with oil. This enables duplex digital protein colorimetric assay with beta galactosidase and alkaline phosphatase enzymes. The platform achieves 5 logs of dynamic range with a limit of detection of 10 aM for both Biotinylated β-galactosidase (B-βG) and Biotin Alkaline Phosphatase Conjugated (B-ALP) proteins. We demonstrate its potential by showing that a higher dosage of paclitaxel suppresses EpCAM-positive EVs but not GPC-1 positive EVs from breast cancer cells, a decline in chemo-resistance that cannot be detected with Western blot analysis of cell lysate. The Mem-dELISA is poised to empower researchers to conduct ultrasensitive, high throughput protein colocalization studies for disease diagnostics, treatment monitoring and biomarker discovery.

## 1. Introduction

Breast cancer is one of the leading malignancies among women in the United States. It ranks second in terms of cancer-related mortality. Despite recent advances in diagnostics and therapy monitoring, such as mammography, magnetic resonance imaging (MRI), and whole-breast ultrasonography, high false positive results limit their effectiveness(Kelly et al., 2010; Roganovic et al., 2015). Moreover, invasive procedures such as tissue biopsies suffer from extended turnaround times, operator bias, and potential misinterpretations owing to tumor heterogeneity, thus making them incompatible for tumor monitoring during treatment with anticancer drugs (Alba-Bernal et al., 2020). In contrast, liquid biopsy offers a promising alternative by analyzing a wide array of biomarkers present in biofluids like blood, saliva, or urine (Connal et al., 2023). It provides insights into the disease status, with minimal invasiveness, comparatively modest cost, and predisposed for high throughput. These molecular biomarkers are often carried by nanocarriers like lipoproteins and Extracellular Vesicles (EVs). Hence, multiplex detection of different colocalized molecules that are markers of tissues of origin, cancer, drug-resistance, or metastasis on the same nanocarriers will provide more specific information about the disease. Therefore, rapid advancements have been made in quantifying several colocalized biomarkers on individual nanocarriers, like lipoproteins and tumor-derived EVs, present in a small volume of physiological fluids. These advances hold promise for early disease diagnostics, prognosis, and therapy management in patients (Cohen and Walt, 2019; Hinestrosa et al., 2022; Kelley et al., 2014; Landegren and Hammond, 2021; Rissin et al., 2010; Wu et al., 2020).

The current state of the art technique for protein detection is Enzyme-Linked Immunosorbent Assay (ELISA) which suffers from insufficient limit of detection (∼nM-pM) beyond which several clinically relevant protein biomarkers (∼pM-fM) of cancer, neurodegenerative, cardiovascular, inflammatory, and autoimmune diseases remain undetected(Wu et al., 2022). To resolve this issue, single-molecule arrays (Simoa) have been developed in which a protein molecule is sandwiched between an antibody coated magnetic beads and an enzyme conjugated detection antibody to form an immunocomplex (Chang et al., 2012; Rivnak et al., 2015). Subsequently, the immunocomplex is sampled into thousands of femtoliter (fL) reaction chambers keeping the total protein concentration within the Poisson limit such that each bead contains either one or zero molecules. The fluorescence signal generated from the enzymatic reaction gives a digital readout providing an absolute quantification of the protein biomarker. This digital ‘on’ and ‘off’ readout format circumvents any bias that usually occurs in other analog sensors based on absolute intensity, current or voltage measurements. For digital quantification of proteins, the focus has been on the fabrication of small microwell-based picoliter reactors, as the enzymatic reaction in ELISA has a linear amplification rate. The ultrasmall volume of a reactor enhances the localized concentration of fluorescent products which can be subsequently detected by a standard fluorescence microscope. Several other methods have been reported in the literature apart from Simoa for digital assay (Rondelez et al., 2005; Sakakihara et al., 2010; Shim et al., 2013; Witters et al., 2013).

However, these digital protein assays are often reliant on specially designed microwells and require complex and expensive cleanroom-based microfabrication techniques. The total number of microwells present in the device determines the sensor’s dynamic range. This is limited by the master mold, which is cumbersome to tune and limits the use of these biosensors for massive multiplexing applications. Moreover, the assays typically involve costly and bulky fluidics control equipment’s such as syringe pumps, centrifuges, valves etc. Additionally, another recurring challenge involves the entrapment of air bubbles within the microwells, potentially impeding precise quantification. Tackling this issue often requires the use of bulky vacuum pumps and costly surface modifications (Zandi Shafagh et al., 2019). Simplifying digital protein assays by reducing the reliance on specialized instruments and mitigating common challenges like air bubble trapping is essential for boarder adoption of the technique in various biomedical applications. Lastly and most importantly, Simoa-based multiplex assays have consistently employed color coding of magnetic beads and utilization of a single enzyme (beta-galactosidase) amplification reaction, which results in maximum of 3-plex immunoassay in a single experimental run (Rissin et al., 2013; Wilson et al., 2016). The absence of multi-color digital ELISA enzymatic amplification makes this assay unsuitable for applications involving protein colocalization assay on EVs, lipoproteins, viruses and proteins having multiple epitopes where the goal is to analyze multiple protein signatures on a single EV captured in a single microwell. Understandably, reports analyzing multiple proteins on EVs using digital ELISA have been performed by repeatedly changing the reporter antibody one at a time(Morasso et al., 2022; Wei et al., 2020).

To address the aforementioned challenges, we report a lithography-free, user-friendly and inexpensive Mem-dELISA digital protein detection platform. The platform utilizes low-cost (<$0.1 material cost) disposable track-etched polycarbonate (PCTE) membranes. The membranes allow reagent loading by wicking through their pores (∼5 µm) before they are adhered to a sticky surface to form thousands of picolitre microwells without generating any air bubble. The Mem-dELISA platform was optimized to perform dual color digital enzymatic reaction of free-floating beta galactosidase and alkaline phosphatase enzymes simultaneously. Magnetic beads were utilized to efficiently capture free floating protein molecules from the sample, with bead loading efficiency into the microwells optimized (>80%) with a permanent magnet and mechanical shaking. The Mem-dELISA digital protein detection biosensor obtained 5 logs of dynamic range (1 pM to 10 aM) for both beta-galactosidase and alkaline phosphate enzymatic amplification with a limit of detection of 10 aM. As a proof-of-concept demonstration, we employed the digital biosensor to perform GPC-1 and EpCAM protein colocalization studies on EVs derived from paclitaxel drug-treated triple-negative breast cancer cell lines (MDA-MB-231 and MDA-MB-468). The study shows that increasing paclitaxel drug dosages resulted in a decrease in the fraction of EVs with colocalized EpCAM-GPC-1 and a decrease in the fraction of EVs expressing EpCAM compared to untreated controls. These trends are not detectable from Western blots of cell lysate. This result clearly suggests the loss of chemo-resistance of cancer cells at high drug dosage, as a decrease in EpCAM expression on EVs has been correlated to loss of metastases and increased survival following drug treatment (Ali et al., 2019; Liu et al., 2019). Additionally, we believe that the user-friendly Mem-dELISA platform’s facile setup, clean room fabrication free microwells formation, low consumables cost (< $0.1) as compared to a typical microfluidics chips (∼ $10), automatic air bubble removal, extremely low instrument cost (compared to > $100k of commercial digital ELISA platforms) and simplified workflows does not require extensive amount of expertise which present a bottleneck for using them for point-of-care applications in resource limited areas where usually a disposable sensor is preferred and expertise may be limited. This platform holds promise for advancing protein detection technologies and improving diagnostics and therapy monitoring in various biomedical applications.

## 2. Materials and Methods

### 2.1 Reagents and materials

Biotinylated β-galactosidase (B-βG) and Biotin Alkaline Phosphatase Conjugated (B-ALP) were purchased from Rockland Immunochemicals (PA, USA). Dynabeads M-280 Streptavidin, Dynabeads™, Biotin conjugated CD326 (EpCAM) monoclonal antibody, RIPA buffer and Eppendorf™ LoBind microcentrifuge tubes were purchased from Thermo Fisher Scientific (MA, USA). Track-etched polycarbonate membranes were purchased from Sigma Aldrich (St. Louis, MO, USA) and Sterlitech Corporation (WA, USA). Resorufin β-D-Galactopyranoside (RDG), Fluorescein di(β-D-galactopyranoside) (FDG), streptavidin-β-galactosidase (S-βG), Biotin Alkaline Phosphatase Conjugated (S-ALP), Tween-20, Bovine Serum Albumin (BSA) and silicone oil were purchased from Sigma Aldrich (St. Louis, MO, USA). The polydimethylsiloxane and curing agent (Sylgard 184 silicon elastomer kit) was purchased from Dow Corning (MI, USA). 4-Methylumbelliferyl phosphate (4-MUP) liquid substrate was purchased from Millipore sigma (St. Louis, USA). Human Glypican 1 (GPC-1) biotinylated antibody was purchased from R&D systems (MN, USA). Biotinylated anti-CD63, anti-vinculin and anti-GPC-1 antibodies were purchased from Abcam (MA, USA). Anti-EpCAM antibody was purchased from Invitrogen (MA, USA) and anti-CD63 was purchased from BD Biosciences (NJ, USA). Anti-rabbit and anti-mouse HRP-conjugated IgG secondary antibodies were purchased from Cell Signaling Technology (MA, USA). Laemmli sample buffer, nonfat dry milk (NFDM) and ECL substrate kit were purchased from Bio-rad (CA, USA). Dulbecco’s Modified Eagle Medium (DMEM) and penicillin-streptomycin were purchased from Corning (NY, USA), and fetal bovine serum (FBS) was purchased from Gibco (NY, USA). Paclitaxel was purchased from Selleck Chemicals (TX, USA).

### 2.2 Device fabrication

A mixture of the curing agent and polydimethylsiloxane (PDMS, Sylgard 184, Dow Corning) base, with a weight ratio of 1:10, was prepared and degassed for one hour to eliminate air bubbles(Yadav et al., 2020). Subsequently, the PDMS was spin-coated onto a 50 x 75 mm glass slide and left overnight for curing, forming a thin film of approximately 200µm. For the creation of the negative mold for the top channel, a 0.3 mm thick piece of KAPTON® Tape (McMaster-Carr) was cut into the shape of a converging-diverging microchannel using a Graphtec Cutting Pro FC7000MK2-60 cutting machine. The tape was then affixed to a petri dish. Then the PDMS was poured into the mold and degassed again to remove any trapped air bubbles, followed by overnight curing at 60°C. After curing, a 1mm biopsy punch was used to punch holes for the inlet and an outlet.

### 2.3 Device assembly

The PDMS thin film was washed with Isopropyl alcohol and then dried with nitrogen gas. A 500 µl drop of 1xPBS was put in the center of the glass slide having a thin film of the PDMS. Subsequently, a piece of PCTE membrane measuring ∼ 1 cm x 0.5 cm was cut and slowly immersed into the droplet using forceps. The membrane was gently pushed until it made contact with the PDMS surface. Finally, the flat surface of the forceps was utilized to ensure the conformal binding of the membrane with the PDMS surface. The membrane demonstrates adhesive characteristics to PDMS, leading to a conformal coating on the elastomeric substrate. The previously fabricated PDMS-based microchannel was placed on top of the membrane and was pressed until it established contact with the PDMS thin film. The excessive buffer solution at the edges of the PDMS channel was then slowly removed using Kimwipes™. The assembled device is shown in Supplementary Figure 1.

### 2.4 Microwell filling experiment using fluorescein

Following the assembly of the device, a 200 µl solution of fluorescein (1mM) was introduced through a pipette into the inlet of the upper PDMS channel. Subsequently, 200 µl of silicone oil was pipetted until it entirely displaced the fluorescein solution in the microchannel. For visualization of the microwell filling performance, MetaMorph v7.7.9 software was used to do z-stack imaging using a Nikon Eclipse Ti confocal microscope attached to an iXonEM+ cooled CCD camera attached to it.

### 2.5 Digital enzyme assay(s) without magnetic beads

After the completion of device assembly steps, a solution comprising 100 µl of B-ALP or/and B-βG (spiked in incubation buffer) each at different concentration were mixed and injected into the microchannel. The solution was incubated for 5 minutes to enable stochastic encapsulation of enzyme molecules inside the microwell. Afterwards, a solution containing 100 µl of 200 μM Fluorescein di(β-D-galactopyranoside) (FDG) and 100 µl of undiluted 4-Methylumbelliferyl phosphate (4-MUP) substrates was prepared and injected into the microchannel. Immediately afterwards, 200 µl of silicone oil was pipetted into the channel to seal all the microwells. The microwells were then imaged after 30 minutes following incubation of the device in a dark chamber. Leica DMi8 inverted microscope was used to image all enzymatic amplification reactions. The DAPI filter was used to measure alkaline phosphate amplification at 1s exposure while amplification of beta galactosidase was observed using Rhodamine filter at an exposure of 500 ms for all imaging steps in digital assay.

### 2.6 Digital enzyme assay(s) with magnetic beads

10 ml stock solutions of washing buffer (5× PBS + 0.1% Tween 20) and incubation buffer (1xPBS + 2% BSA + 0.05% Tween 20) were prepared and stored at 4 °C for subsequent use. 50 µl of magnetic beads (Dynabeads M-280 Streptavidin, Dynabeads™) from the commercial vial (10 mg/ml) was taken and diluted 25 times to make a stock solution of magnetic beads. Subsequently, a 50 µl of this stock solution was taken in low protein binding microcentrifuge tubes, washed 3 times with washing buffer and incubated in the incubation buffer for a duration of 2 hours. Following the incubation, the beads were washed 3 times and were subsequently incubated with 100 µl of incubation buffer spiked with B-βG and/or B-ALP into the tube. This mixture underwent a 2-hour incubation period while being rotated. Afterwards, the tubes were washed 3 times in the washing buffer to remove nonspecifically bound molecules and incubated in the incubation buffer to a final volume of 100 µl. After wetting the membrane as outlined in ‘Device assembly’ section, the solution containing magnetic beads was pipetted on top of the wetted membrane. The NdFeB cylinder magnet (DCC-N52, K&J Magnetics) was kept at an optimized distance below the PDMS coated glass slide. The glass slide was manually shaked in the xy plane for 15 minutes to enhance bead filling within the microwells. Later, the PDMS based microchannel was gently put on the top to complete the device assembly as described earlier. Subsequently, a solution consisting of 100 µl of 200 μM FDG and 100 µl of undiluted 4-MUP substrates were mixed in PBS and introduced into the microchannel. Immediately thereafter, 200 µl of silicone oil was pipetted into the channel to achieve the sealing of all microwells. The device was then incubated in a dark chamber and was subsequently imaged after half an hour.

### 2.7 Magnetic bead functionalization with antibodies

The 50 µl stock solution of streptavidin conjugated magnetic beads was aliquoted in low protein binding microcentrifuge tubes, washed 3 times with washing buffer and incubated in the incubation buffer for a duration of 2 hours. 50 µl of 100x (diluted in PBS + 0.05% Tween 20) diluted Biotin-conjugated CD63 antibody were incubated with the magnetic beads in a shaker for 3 hours. Following this incubation, the beads underwent three successive washes in the washing buffer and were subsequently stored in the incubation buffer at 4 °C until further use.

### 2.8 Extracellular Vesicles (EVs) isolation

MDA-MB-231 and MDA-MB-468 cells were cultured in DMEM supplemented with 10% FBS and 1% penicillin-streptomycin at 37°C with 5% CO_2_. EV-depleted media was made with FBS that had been centrifuged at 100,000 x g for 9 hours to remove bovine EVs. EV-depleted media was added to confluent flasks and incubated for 48 hours. For paclitaxel treatment conditions, paclitaxel was added to the EV-depleted media. The media was then collected and spun at 400 x g for 5 min to remove cell debris, and the supernatant was centrifuged at 100,000 x g for 1 h to pellet EVs. The EV pellet was washed once with PBS, then resuspended in PBS and stored at -80°C for downstream analysis.

### 2.9 Multiplex digital protein assay of EVs

The complete workflow of digital ELISA is described in Figure 1C. The 50 µl of CD 63 antibody functionalized magnetic beads described earlier were incubated with 100 µl of EVs (100x diluted in incubation buffer from ultracentrifugation) for 2 hours under constant rotation. The beads were washed 3 times in the washing buffer and were later incubated with the incubation buffer. Parallelly, biotinylated GPC-1 detection antibody was incubated with 20 pM of S-ALP and 1 nM biotinylated EpCAM was incubated with 20 pM of S-βG respectively for 2 hours to attach enzymes on antibodies. The stored magnetic beads were then incubated with 100 µl of GPC-1 & S-ALP enzyme complex for 30 minutes under rotation. The immunocomplex was then washed 8 times with the washing buffer to remove non-specifically bound molecules. Similarly, 100 µl of biotinylated EpCAM & S-βG complex was then incubated with the beads for half an hour under constant rotation and was washed 8 times to form the immunocomplex. This magnetic bead based immunocomplex was used to perform digital protein detection using the steps described in earlier sections.

**Figure 1:**
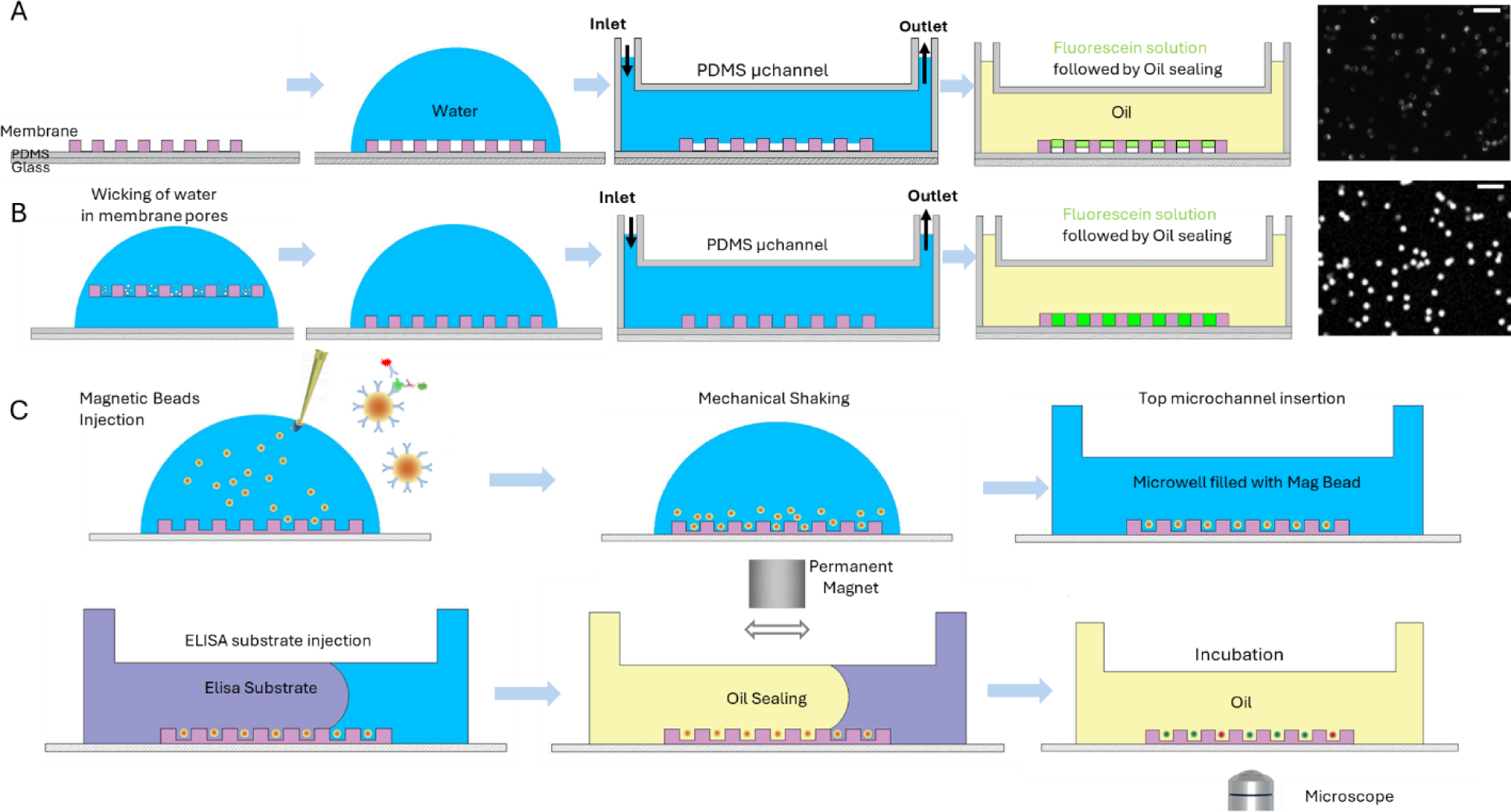
Schematics of comparison of two workflows to use the through holes of the PCTE membrane to obtain microwells for digital assay. **A)** The membrane is first conformally sealed on the PDMS layer and then wetted with water. After inserting a PDMS micro channel from the top, fluorescein solution was inserted followed by oil sealing. The fluorescence images show trapped bubbles inside microwells. **B)** The membrane is first wetted with water to remove all trapped air bubbles due to wicking. Afterwards, the membrane is conformally assembled over the PDMS layer and a PDMS micro channel is inserted from the top. The fluorescence images obtained by first putting the fluorescein solution and then followed by oil sealing show the microwells are filled without trapped bubbles. Scale bar is 20 µm. **C)** Mem-dELISA device workflow for utilizing the through holes of the membrane for digital ELISA.

### 2.10 Western Blot

Cells were lysed in RIPA buffer (Thermo Scientific) and spun at 16,000 x g for 20 min to obtain the protein supernatant. Samples were prepared for SDS-PAGE in Laemmli sample buffer (BIO-RAD) and heated to 70 °C for 10 minutes. SDS-PAGE was performed with 20 ug of protein per sample in a polyacrylamide gel, and the protein was electrophoretically transferred to a nitrocellulose membrane. The membranes were blocked with 5% nonfat dry milk (NFDM, BIO-RAD) TBS with Tween-20 (TBST, abcam), then incubated overnight with a primary antibody mixture containing either anti-vinculin (1:5000, abcam), anti-GPC-1 (1:1000, abcam) and anti-EpCAM (1:50, Invitrogen) primary antibodies or anti-vinculin and anti-CD63 (1:500, BD Biosciences) antibodies in TBST with 5% NFDM. After washing with TBST, the membranes were incubated for 1 hour in TBST containing anti-rabbit (1:1000, Cell Signaling Technology) and anti-mouse (1:1000, Cell Signaling Technology) HRP-conjugated IgG secondary antibodies. After washing with TBST, the protein was visualized using an ECL substrate kit (BIO-RAD) and imaged with a ChemiDoc-It2 system.

### 2.11 Image processing and data analysis

All fluorescence images were processed using a custom developed code. More details can be found in Supplementary Figure 2. GraphPad Prism has been used for graphical representation along with statistical analysis. Unless otherwise specified, all data used in this study are shown as mean ± standard deviation.

## 3. Results and discussion

### 3.1 Membrane as microwells chamber

PCTE membranes, commercially produced through ion irradiation and subsequent track etching, exhibit a high density of uniformly sized cylindrical through-holes. These holes are available in a range of desired diameters (from 10 nm to 30 μm) and possess low protein binding properties (Dutt et al., 2021; Lin et al., 2019). In the Mem-dELISA technology, we ingeniously utilize the through holes of the PCTE membranes to repurpose them as microwells by strategically blocking one end as described in Figure 1. To accomplish this, a drop of 1xPBS was first put on top of a glass slide spin coated (1000 rpm, 2 minutes) with a thin film of PDMS (Figure 1B). The membrane was fully immersed into the PBS solution using forceps to remove all the air bubbles via capillary wicking of water inside the through holes. After successfully eliminated all trapped air bubbles, we capitalize on the adhesive nature of the PCTE membrane to a PDMS surface. The forceps were gently used to place the membrane on the PDMS surface, where it adhered to form microwells. Afterwards, a PDMS based microchannel was put on top to develop the integrated Mem-dELISA platform. The PDMS based microchannel permits the flow of different reagents on top of membrane, diffusion of reagents to the microwells and ensures complete sample partition within each microwell after sealing with oil. Injection of a fluorescein solution followed by oil sealing showed complete segregation of microwells by oil as shown in the fluorescent image obtained in Figure 1B. The initial wicking action of the membrane is crucial to the removal of trapped bubbles. If the membrane is first attached to the PDMS surface followed by the placement of the PBS droplet above, trapped air bubbles were consistently observed in the majority of microwells (Figure 1A). Using commercially available micron-sized (∼2.8 µm) magnetic beads for protein capture, we opted to develop our Mem-dELISA technology by utilizing a 5 μm pore size membrane. The selected membrane exhibited uniformly sized microwells with a low population of overlapping pores as evident from the bright field image (see Supplementary Figure 3).

With the initial workflow finalized, we optimized the height of the PDMS based microchannel to enable seamless sealing of microwells by oil (Figure 2A). After the device assembly, a 1 mM solution of fluorescein was injected into the microchannel followed by oil injection to partition individual microwells. Two distinct types of hydrophobic and hydrophilic commercial membranes were also utilized in the experiment to understand the effect of the surface property of membrane. The height of the microchannel was systematically varied for each type of membrane from 100-1000 µm. Since the surface of the PCTE membrane is uneven, a high shear is necessary to remove the remnant water layer present on the top of the membrane. For hydrophilic membrane, the layer of water could not be removed until the microchannel height was reduced to 100 µm. Even with this lowered channel height, many water islands were observed thus corrupting sample partitioning (Figure 2C). In case of hydrophobic membranes, excellent sample partition was observed when microchannel height was reduced to 300 µm and less as shown in Figure 2B and 2D. The images show a clear separation of individual picolitre reactors containing a fluorescein solution. From these findings, we selected the microchannel height of 300 µm for our subsequent experiments. Z-stack confocal imaging was performed to orthogonally confirm the removal of all air bubbles within the microwells in the hydrophobic membrane as shown in Figure 2E. As we move in z-direction, the microwells transition into focus and recede out of focus at 25 µm. The results indicate that the height of the cylindrical pores is around 20 µm which aligns closely to the manufacturers specifications of 21 µm. Using these results, we estimate that the volume of one microwell is ∼0.4 pL while the membrane has a well density of 4 × 10^5^ *wells/cm*^2^.

**Figure 2:**
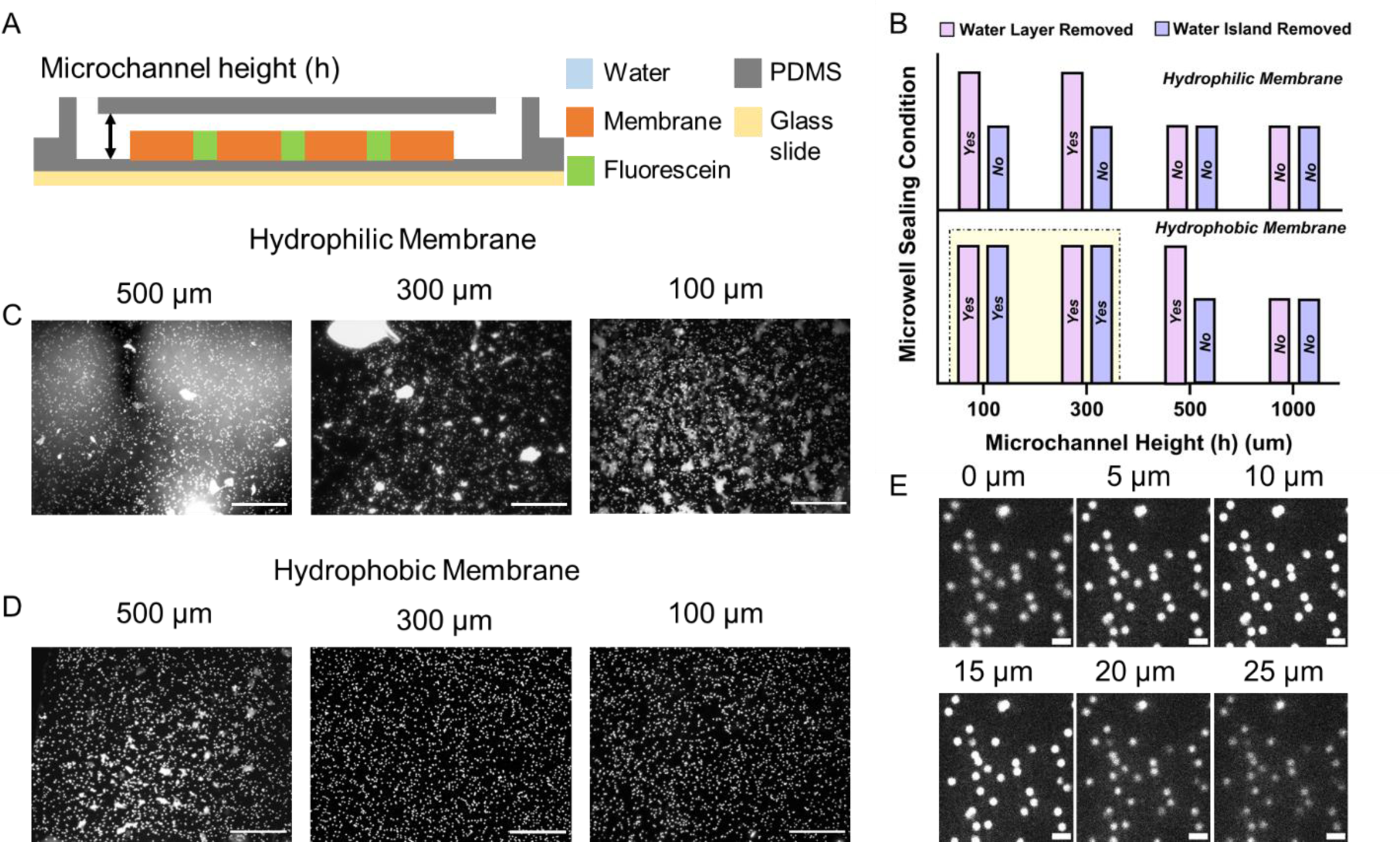
Optimization of micro-channel height of Mem-dELISA device for perfect sealing of microwells. **A)** Schematics showing the height of the PDMS micro-channel used for injecting fluorescent liquid and oil for sealing of microwells. **B)** Graph depicting the parametric variation of micro-channel height to study its effect on the removal of fluorescent liquid by oil. The oil sealing method was very effective to remove the remnant water layer in hydrophobic membrane at a gap height of 300 µm and 100 µm. **C & D)** Images depicting the sealing of microwells by oil after filling them with fluorescent solution for both hydrophilic and hydrophobic PCTE membranes. **E)** Confocal z-stack images of microwells filled with 1mM fluorescein solution and sealed with oil depicting no crosstalk between individual microwells. Scale bar is 200 µm for C) & D) and 10 µm in E).

Since we are utilizing the wicking to remove all trapped bubbles inside the membrane, our method provides an extremely low-cost solution for microwell fabrication and operation as compared to other methods that utilize complex lithography techniques. Additionally, it eliminates the need for expensive vacuum pumps and surface modification of microwells to render them super-hydrophilic. Furthermore, the quantity of microwells is contingent upon the size of the membrane and can be readily scaled to ∼10^6^ wells with a ∼2.5 *cm*^2^ section of 5 µm pore size membrane for highly multiplex assays. This distinguishes our technique from other fabrication-based methods where the microwell count is constrained by the dimensions of the master mold.

### 3.2 Single-molecule assay for free floating B-βG and B-ALP

Since the commercialization of Quanterix’s Simoa® technology for digital ELISA, great progress has been made in utilizing the power of digital counting of protein molecules using femtoliter-sized chambers. Notably, the enzymatic amplification in their device is executed exclusively through the utilization of beta-galactosidase enzymatic reaction with its substrate. It is imperative to recognize that a solitary enzymatic amplification within a singular microwell is insufficient for elucidating the colocalization of multiple proteins within nanocarriers, such as extracellular vesicles which are extensively studied nowadays.

In this study, we have selected beta galactosidase and alkaline phosphatase as our two candidate enzymes for performing duplex enzymatic reaction within the same microwell. To perform duplex assay, it’s important to test the enzyme cross reactivity with their substrates (Obayashi et al., 2015; Ono et al., 2018). Various enzyme-substrate combinations were systematically tested to ascertain the combination exhibiting minimal fluorescence crosstalk and inhibitory behavior as summarized in Supplementary Figure 4. From our study, beta galactosidase-RDG and alkaline phosphatase-4 MUP substrate combination showed the best results. The B-βG reacts with RDG to give red fluorescence signal (resorufin: Emission 584 nm) signal and B-ALP reacts with 4-MUP to give a blue fluorescence signal (methylumbelliferone: Emission 445 nm) that can be separated by using Rhodamine and DAPI filters respectively. We also performed a bulk cross reactivity study in which the concentration of B-βG varied from 0 to 20 nM while keeping the B-ALP concentration fixed at 0.7 nM (Supplementary Figure 5). The blue fluorescence remained roughly constant while the green fluorescence increased with the concentration of B-βG confirming minimum cross reactivity between the two enzymes.

After successful demonstration of bubble-free partitioning of liquid into microwells, the Mem-dELISA platform was tested to study single molecule amplification of B-βG and B-ALP enzymes. The complete workflow is described in Supplementary Figure 6. Following the device assembly and filling each microwell with PBS, a solution of B-ALP (300 pM) was quickly injected into the chip. The chip was then incubated for a minute, allowing for the stochastic encapsulation of B-ALP molecules within the microwells through diffusion from the microchannel to the microwells. Afterwards, 2x diluted 4-MUP substrate was injected into the microchannel followed by sealing of the microwells with oil. The time elapsed images of these microwells are shown in Figure 3A where the fluorescence intensity increases with time depicting enzymatic reaction between B-ALP and 4-MUP substrate. The intensity of each microwell at 25 minutes image was extracted using a custom developed code and plotted as a histogram, fitted with the sum of four Gaussian functions depicting the presence of 0, 1, 2 and 3+ enzymes in the microwells (Figure 3B). Theoretically, the encapsulation of enzyme molecules in the empty microwells follows a Poisson distribution: 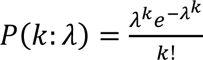. Here *P*(*k*: *λ*) denotes the random probability of encapsulating k molecules in a microwell and *λ* is the average number of molecules per microwell. At the calculated lambda of 0.42, the expected probabilities of 0, 1, 2 and 3+ enzymes per microwells are 65.71%, 27.60%, 5.80% and 0.01% respectively. The experimentally observed values obtained from Gaussian fitting matched well with the theoretical estimates from Poisson distribution as shown in Figure 3C. The presence of discrete peaks is consistent with earlier reports (Obayashi et al., 2015). Similarly, we did the same experiment to study the enzymatic amplification of beta galactosidase enzyme with its substrate (100 μM RDG) by keeping B-βG enzyme concentration at 30 pM. The sequential images capturing the temporal evolution of fluorescence intensity increase in microwells are presented in Figure 3D. As described earlier, the fluorescence intensity associated with each microwell was extracted from the image captured at the 25 minutes timestamp. Subsequently, a histogram was generated and superimposed with the composite fit derived from the summation of four Gaussian functions (Figure 3E). The obtained experimental values matched well with the theoretical estimates from Poisson distribution with *λ* = 0.224 (Figure 3F). The discrete fluorescence peaks observed with B-βG enzyme and RDG substrate reaction in microwells are also consistent with the previous literature reports (Rondelez et al., 2005; Sakakihara et al., 2010).

**Figure 3:**
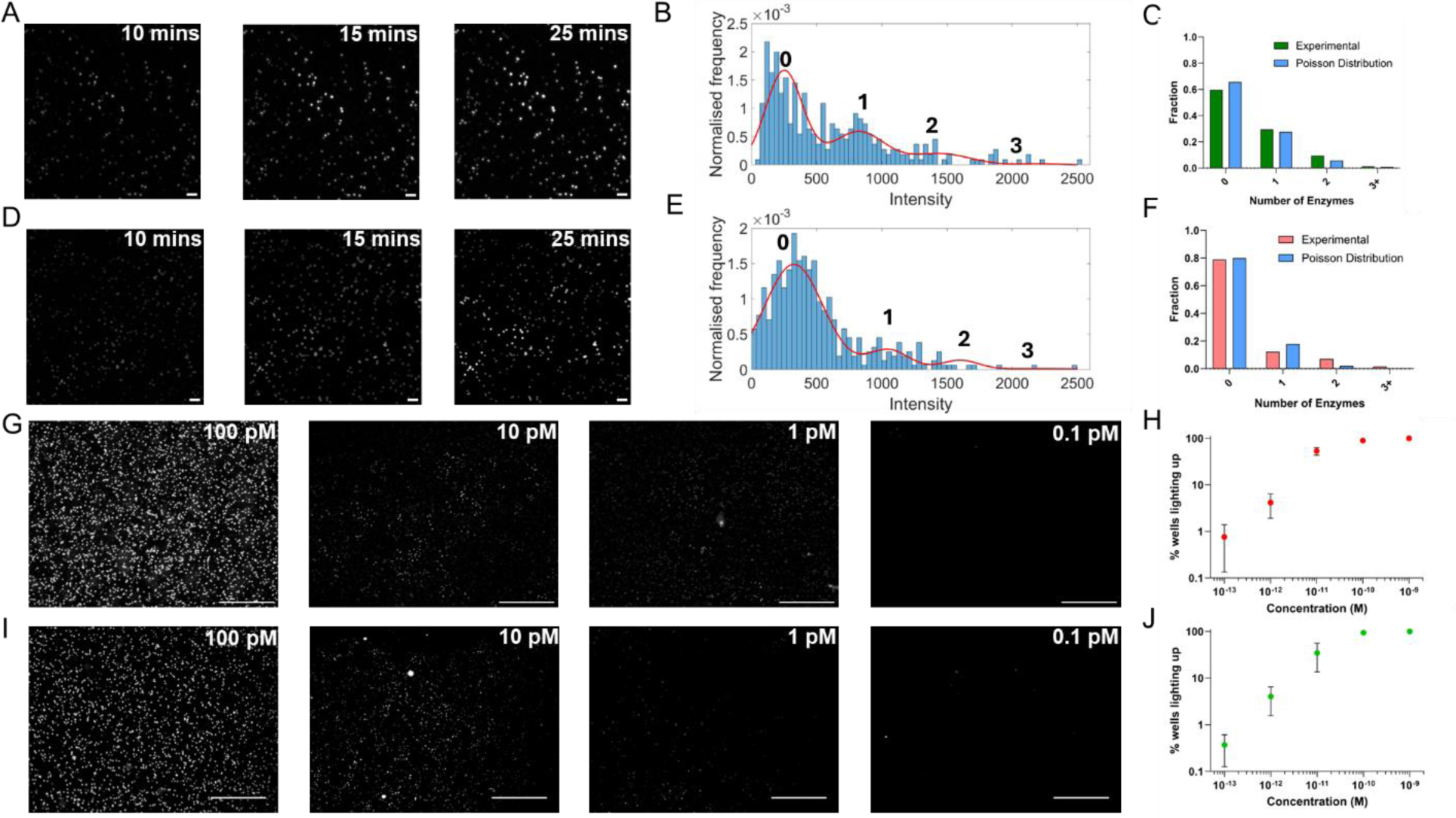
Single molecule amplification results of B-ALP and B-βG in Mem-dELISA device. **A)** Sequential fluorescent images for digital enzymatic amplification for B-ALP at 10,15 and 25 mins. **B)** The intensity distribution of each microwell extracted from the 25 mins image and fitted with sum of four gaussians corresponding to microwells having 0,1,2 and 3+ B-ALP molecules. **C)** Comparison of the % of microwells having 0,1,2 and 3+ B-ALP molecules obtained experimentally by gaussian fitting and obtained by Poisson’s distribution having *λ* of 0.42. **D)** Temporal fluorescent images for digital enzymatic amplification for B-βG at 10,15 and 25 mins. **E)** The intensity distribution of each microwell extracted from the 25 mins B-βG image and fitted with sum of four gaussians corresponding to microwells having 0,1,2 and 3 B-βG molecules. **F)** Comparison of the % of microwells having 0,1,2 and 3 B-ALP molecules obtained experimentally by gaussian fitting and obtained by Poisson’s distribution having *λ* of 0.224. **G)** Representative fluorescence images of serial concentration dilution (100 pM – 0.1 pM) of B-βG. **H)** The corresponding log-log plot of the % active wells as a function of B-βG protein concentrations (n=3). **I)** Illustrative fluorescence images of serial dilution in concentration (100 pM – 0.1 pM) of B-ALP molecules and **J)** the corresponding plot of % active wells vs B-ALP concentration. Scale bar is 50 µm for A) & D) and 200 µm in G) & I).

The analysis shown above confirms the encapsulation and amplification of single molecules of B-βG and B-ALP within individual microwells in separate experiments at a particular concentration. We now proceeded to vary the concentrations of both B-βG and B-ALP in separate experiments within the range of 100-0.1 pM. With the sequential reduction in enzyme concentrations, the total number of wells lighting up decreased exponentially based on Poisson statistics as shown in the series of fluorescent images in Figure 3G and 3I. A log-log plot revealing the percentage of fluorescent microwells as a function of the concentration is depicted in Figure 3H and 3I. The biosensor had a working concentration range 10-0.1 pM for both B-βG and B-ALP. This analysis shows that the Mem-dELISA methods and protocols have no effect in the biochemical reaction between different enzymes and their respective substrates.

After optimizing the individual reactions of B-βG and B-ALP separately on the microwells, duplex assay was performed. The B-βG (30 pM) and B-ALP (100 pM) were mixed and incubated in the microchannel for a minute. The RDG and 4-MUP substrates were mixed and immediately injected into the microchannel followed by immediate sealing with oil. It is important to note that the final temperature of the mixture influenced the reaction kinetics (data not shown). In our experience, both the substrates are warmed up to 37 °C before mixing them together for optimal results. Supplementary Figure 7 displays a superimposed fluorescence image derived from the blue and red fluorescence channels. It is important to note that for better visualization of overlayed images, the blue channel images have been colored with green hues. Therefore, the green and red hues correspond to the fluorescence signature of 4-MU and resorufin, respectively. Consistent with our expectations, the green and red fluorescent channels exhibited a random distribution, with certain microwells displaying both green and red fluorescence concurrently (yellow/orange microwells) which indicates co encapsulation of both B-βG and B-ALP in the same microwell. Hence, successful demonstration of the feasibility of dual-color digital enzyme assays is presented in this section and will be explored further in the upcoming sections.

### 3.3 Magnetic beads seeding optimization

In the earlier section we demonstrated the capture of free-floating beta galactosidase and alkaline phosphate enzyme molecules in microwells for performing digital assays. However, relying only on diffusion to capture free-floating protein particles leads to detection of a very small fraction of protein molecules as more than 99.9% of the sample is lost. Hence, to increase the sensitivity of the digital sensor, magnetic beads are used to concentrate the free-floating proteins into the microwells (Lim and Zhang, 2007). Moreover, to increase the percentage of magnetic bead inside the microwells, permanent magnets have been used although it can cause adverse effects such as bead chain formation that significantly reduces the bead capture efficiency (Verbruggen et al., 2015). To develop more mechanistic insights of the system, COMSOL Multiphysics simulations of a permanent bar magnet was performed using the ‘Magnetic fields, no current’ module to understand the variation of magnetic flux density(Sharma et al., 2018) as a function of distance from the permanent magnet as shown in Figure 4A. The simulation geometry and details are presented in Supplementary Figure 8. The simulations show that the magnetic flux density varies as ∼1/*r*^3^ with r is the radial distance from the magnet Figure 4B. Moreover, the magnetic force exerted on the magnetic bead by the permanent magnet is directly proportional to the square of the gradient of the magnetic field. This relationship also underscores the sensitivity of the force acting on the magnetic bead to variations in radial distance from the permanent magnet. Unfavorably, the strong field from the magnet often causes undesirable chain formation between magnetic beads which negatively impacts the digital assay. Hence, the distance of the membrane surface from the permanent magnet needs to be optimized so to concentrate the beads on the membrane surface without causing undesirable chain formation.

**Figure 4:**
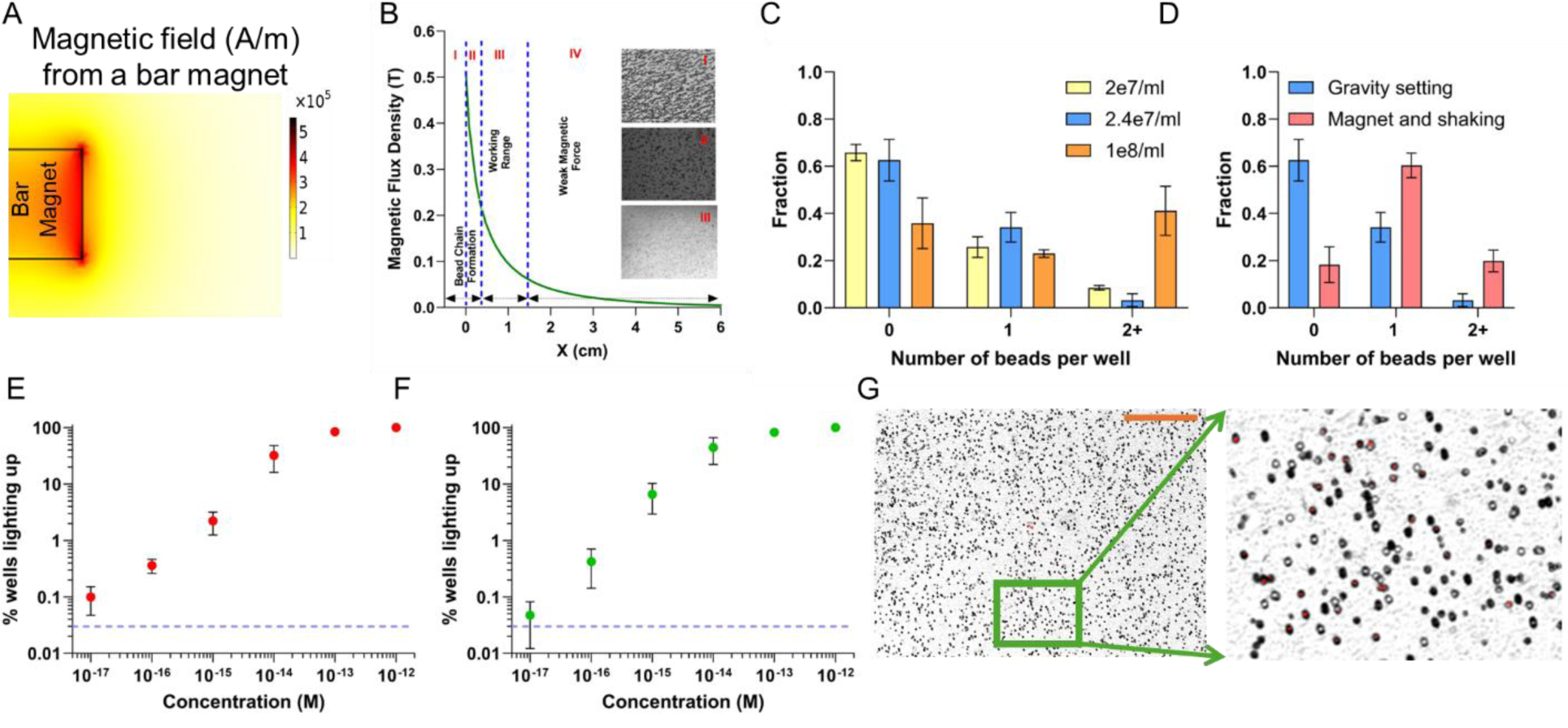
Digital protein detection of B-ALP and B-βG protein molecules using magnetic beads in Mem-dELISA device. **A)** The surface plot of magnetic field from a permanent magnet obtained from COMSOL Multiphysics simulations. **B)** The variation of magnetic flux density (T) as a function of axial distance from the midpoint of the magnet obtained from FEM simulations. The experimentally obtained images of magnetic beads were obtained by varying the permanent magnet distance from the membrane surface. The bead phenotypes are divided into 3 zones. In I and II, magnetic beads chain formation was obtained which significantly reduces bead capture in the microwells. Regime III is the working range whereas in regime IV the magnetic force from the magnet didn’t have any impact on the beads. **C)** The loading efficiency of magnetic beads in microwells without using a magnet for three different magnetic bead densities. **D)** Comparison of magnetic beads loading in microwells with gravity settling and using a magnet with mechanical shaking for 15 minutes at bead density of 2.4e7/ml. **E)** & **F)** A log-log graph of % of fluorescent wells variation with B-βG and B-ALP protein concentration respectively (n=4 minimum). **G)** The overlay of bright field and red fluorescent image. The zoomed image depicts the active wells (red) and inactive wells. Scale bar is 200 µm.

The experimental results indicate the occurrence of large undesired magnetic bead chains when the beads were directly positioned atop the magnet, as illustrated in zone I of Figure 4B. Note that the origin is taken on the plane of the membrane which is ∼1.2 mm distant from the magnet’s surface. With an increasing separation between the magnet and the membrane, the prominent large chunks of bead aggregates progressively diminish as indicated by zone II image. Moreover, the large bead chain formation was greatly reduced when the magnet was positioned approximately 4 mm away from the glass slide. The operational effectiveness of the magnetic force was found to be optimal within the range of 4 mm to 1.6 cm from the glass slide, where no significant formation of magnetic bead chains was observed. Beyond 2 cm, the magnetic force was not sufficient to pull all the free-floating beads in the solution phase on top of the membrane. For all the experiments, the magnet was kept at 5 mm from the glass slide.

Initial experiments were performed with magnetic beads without using a permanent magnet. The magnetic beads themselves are expected to be captured into the microwell following Poisson distribution, imposing a theoretical limit of the maximum percentage (≤ 37%) of single bead occupancy in the microwell chamber (Chang et al., 2012). We observed the same behavior when we seeded the magnetic beads at three different bead densities on top of the membrane piece and utilized gravity settling of beads to fill the microwells (Figure 4C). The majority of the microwells remained empty in these cases with less than 40% of the microwells are filled with magnetic beads by using gravity settling (Supplementary Figure 13). As the bead density was increased to 10^8^*beads*/*ml*, single bead occupancy decreased, and double bead occupancy increased drastically. Hence, 2.4 × 10^7^*beads*/*ml* density was selected for further optimization using a magnet. After the beads settled on the membrane surface, the Mem-dELISA chip was mechanically shaken for 15 minutes under the influence of the magnetic field from a permanent magnet. This technique improved the single bead capture from ∼34% to >60% (Figure 4D). In total, the final bead capture efficiency was increased to > 80% using a permanent magnet and mechanical shaking.

We offer a simple mechanistic explanation for this improved result. During mechanical shaking, an impulse is given to the particle in the horizontal direction while the magnet is kept in the vertical direction. A simple force balance on a magnetic bead reveals that the drag acts in the horizontal direction whereas the magnetic force along with gravity acts in the vertical direction. The net resultant vector will be inclined at an angle to the vertical direction. Hence the bead will bounce on the membrane, thus substantially elevating the probability of locating a nearby microwell. Once the bead is captured into the microwell, the high surface tension prevents the bead from exiting the microwells.

Another notable point is that the microwells in our membranes are more separated and deeper as compared to other devices. During processes such as dendritic growth or chain formation of magnetic beads, the trapped particles amplify the magnetic field, thereby capturing additional particles atop them. Importantly, this enhanced field is limited to only a few bead diameters. Hence, we hypothesize that well-separated wells (beyond one particle diameter) and large number of microwells is the key to prevent chain formation and improve sensitivity. The randomly distributed and well-separated deep microwells of the PCTE membrane facilitate the convenient utilization of a permanent magnet to increase the bead loading efficiency.

### 3.4 Magnetic bead-based multiplex assay

Following the optimization of the magnetic bead loading step, digital protein assays were conducted on the Mem-dELISA device to improve the limit of detection from Pico (10^−12^) to 10 Atto (10^−18^) molar range for B-βG molecules captured by streptavidin-coated magnetic beads. For an initial test, 1 pM concentration of B-βG was incubated with streptavidin-coated magnetic beads. At this concentration, it is expected that each magnetic bead should capture at least one copy of B-βG, if not more. To confirm this hypothesis, both brightfield and its corresponding fluorescence image were taken at 50x resolution for comparison (Supplementary Figure 9). The microwells that contain no magnetic bead are in ‘off’ state, while microwells that contain a magnetic bead are in ‘on’ state (superimposed red color). The red marked circles on the fluorescence image are the empty microwells that did not show any signal. This confirms that only the magnetic beads present in the microwells were in ‘on’ state. Next, the concentration of B-βG was reduced to 20 fM and a similar analysis was performed. At this concentration, only a fraction of magnetic beads should capture B-βG molecule, and a composite image is made using both brightfield and fluorescence (Red) as shown in Figure 4G. The zoomed-in image shows the presence of empty microwells, microwells that contain a magnetic bead and are in ‘off’ state and microwells that contain a magnetic bead and are in ‘on’ state (superimposed red color). These initial tests validated the capture of B-βG by streptavidin-coated magnetic beads.

Hence, the concentration of B-βG was systematically reduced by an order of magnitude from 1 pM to 10 aM as shown by the series of images in Supplementary Figure 10A. Figure 4E shows a log-log graph depicting the variation of % of fluorescent wells as a function of B-βG protein concentration. The dotted line is obtained from the negative controls, showing that 0.03% of microwells will be fluorescent. The calibration curve showed 5 logs of dynamic range (100 fM-10 aM) and limit of detection of ∼10 aM. It is important to note that the utilization of magnetic beads for capturing free-floating protein molecules has resulted in an improved limit of detection by 5 orders of magnitude, shifting from 100 fM to 10 as compared to cases without magnetic beads (Figure 3H). Similarly, the concentration of B-ALP was also varied from 1 pM– 10 aM as shown in Supplementary Figure 10B and a similar calibration plot was made as discussed above in Figure 3F.

Next, we tested the Mem-dELISA device to perform duplex digital protein assay by detecting both B-βG and B-ALP proteins simultaneously in a single experiment (Figure 5A). Since the base substrates used in these reactions (RDG and 4-MUP) also produce a fluorescent signal, it was necessary to perform a negative control to obtain the baseline intensity. The intensities of all microwells were extracted from images (Figure 5B) captured from both fluorescence filters (Blue and Red) and plotted as histograms as shown in Figure 5C and 5D. As observed in negative controls, <0.03% of the microwells light up at an intensity threshold of five standard deviations from the mean value. Consequently, a microwell is designated as being in an ‘on’ state if its intensity exceeds this specified threshold. To perform the duplex digital assay, for the first experiment, 1 pM of B-ALP and 100 fM of B-βG were mixed and incubated with magnetic beads. As expected, at these concentrations, the majority of microwells were in ‘on’ state as shown in Figure 5E. The concentration of both protein molecules was serially decreased by an order of magnitude in subsequent experiments and the results were shown as composite images in Figure 5 (H, K, N & Q). Since the blue fluorescence from 4-MUP substrate was consistently present in all microwells, its fluorescence was used for accurate position identification for all microwells using an in-house developed MATLAB script. As the concentration decreases by an order of magnitude, the number of microwells crossing the threshold intensity decreases as well for both B-βG and B-ALP. For efficient data visualization, the histogram of the intensity of microwells was plotted for all concentrations of B-βG and B-ALP in Figure 5 (F, G, I, J, L, M, O, P, R & S). The threshold intensity in these histograms is obtained from the negative controls described earlier. As the concentration of the protein molecules decreased, the histogram peaks shifted towards the left indicating that the number of microwells lighting up also decreased significantly. Additionally, the microwell intensity from each microwell was also extracted from a representative image and plotted as a function of the microwell ID for various concentrations in Supplementary Figure 11 for better visualization of this trend. Furthermore, the capture of protein molecules by magnetic beads shall also obey the Poisson’s distribution as described in previous reports (Liu et al., 2018). For a duplex assay, since a single bead is present to capture two free floating protein molecules, the combined probability of capturing both protein molecules on the same magnetic bead shall be the multiplication of two independent Poisson distributions. In Figure 5G, the percentage of microwells having both B-βG and B-ALP proteins is plotted as a function of their concentrations which matches well with the theoretical predictions of double Poisson’s distribution.

**Figure 5:**
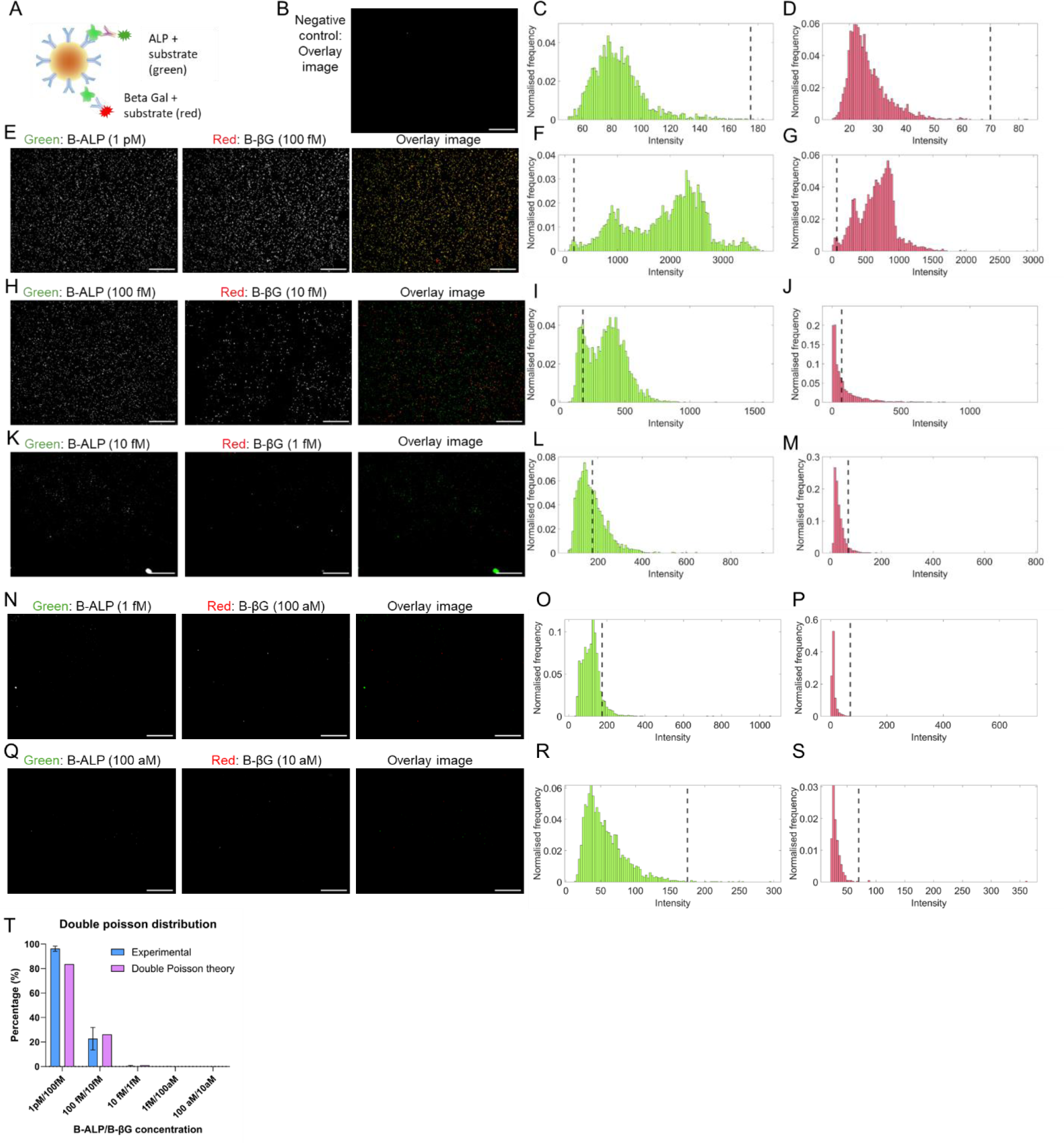
Dual color protein detection using magnetic beads in Mem-dELISA device. **A)** Schematic of dual color protein detection. **B)** Overlay image of the negative control (mixture of 4-MUP (green color) and RDG (red color)). *Note that even though we look at blue fluorescence to observe ALP amplification, the image is shown in green color for better visualization of the overlay image*. **C** & **D)** Histogram of intensity distribution for negative control from both blue fluorescence and red fluorescence channels. The dotted line in both histograms represents the intensity value equal to mean + 5*standard deviation. **E, H, K, N & Q)** Fluorescence images from both filters along with the overlay image for the detection of B-ALP and B-βG simultaneously in the Mem-dELISA device. The concentrations of both B-ALP and B-βG proteins were serially diluted while keeping their ratio constant. The B-ALP concentration varied from 1 pM to 100 aM while B-βG concentration varied from 100 fM to 10 aM. **F, G, I, J, L, M, O, P, R & S)** Histograms of intensity distribution of microwells for both B-ALP (green bins) and B-βG (red bins) amplification in microwells with various concentration as mentioned in **E, H, K, N & Q.** The dotted line intensity was obtained from the negative control histogram. **T)** B-ALP and B-βG colocalization percentage and its comparison with double Poisson’s statistics prediction at various concentrations, mentioned in B, C, D, E, F. Scale bar is 200 µm for all images.

### 3.5 EVs based dELISA

EVs are an emerging class of promising highly heterogeneous circulating biomarkers that play important roles in shuttling molecular cargo from host cell to recipient cell, thereby facilitating intercellular communication, modulating drug resistances and immune response (Schwarzenbach and Gahan, 2020; Wills et al., 2021). Therefore, by first principles, alterations in the protein expression of EVs derived from tumors are expected to exhibit a strong correlation with the protein expression in the host tumor cells. In the case of breast cancer, the molecular cargo of EVs has been associated with prediction of therapy outcome and drug resistance (Ciravolo et al., 2012; Yu et al., 2016). Though the EVs heterogeneity in size and molecular cargo has been well documented, bulk EV analysis methods (Dynamic Light Scattering, Nanoparticle Tracking Analysis, ELISA and Western blots) have been predominantly used while advances have been made to perform single EV analysis using single-particle interferometric reflectance imaging with fluorescence, nanoparticle tracking analysis, microfluidic resistive pulse sensing, and nanoflow cytometry (Arab et al., 2021; Shao et al., 2018; Sharma et al., 2023). Nonetheless, most single EV analysis platforms that analyze the molecular biomarkers suffer from low dynamic range (∼2 logs), interference from non-targets and inherent issues with fluorescent probes such as protein autofluorescence and photobleaching. The developed dual color Mem-dELISA biosensor with 5 logs of dynamic range utilizes enzymatic amplification instead of fluorescent labelling and can easily overcome these challenges. Moreover, the platform utilizes efficient wash protocols using high ionic strength buffer (5X PBS) to minimize electrostatic interactions between non targets and beads. The high throughput analysis of different surface proteins simultaneously on the surface of a single EV without EV lysis provides a holistic approach to capture the heterogeneity with improved reproducibility to allow accurate diagnostic and therapeutic predictions. Multiple protein analysis on the same EVs improves normalization of data with a reference protein to minimize experimental bias caused by upstream EV isolation steps (Sharma et al., 2023).

We utilized the multiplex Mem-dELISA platform to detect the two proteins colocalized on the surface of a single EV derived from cell culture media of breast cancer cell lines. As a proof-of-concept study, we studied the effect of chemotherapy (paclitaxel) treatment on two triple-negative breast cancer cell lines: MDA-MB-231 and MDA-MB-468 by quantifying the colocalization of GPC-1 and EpCAM proteins on the surface of EVs. Several reports suggest that both EpCAM and GPC-1 are biomarkers of breast cancer and are present in both MDA-MB-231 and MDA-MB-468 cell lines (Li et al., 2018; Lu et al., 2021; Matsuda et al., 2001). EpCAM has been attributed to increased drug resistance and poor prognosis (Soysal et al., 2013; Tian et al., 2021). We hypothesized that the normalized ratio of colocalized EpCAM-GPC-1 on EVs would provide insights on the effect of paclitaxel drug treatment on chemo-resistance for use in drug screening and therapy management.

Magnetic beads coated with CD63, a known tetraspanin marker, were selected to capture EVs isolated by ultracentrifugation from cell culture. An immunocomplex was formed where an EV is sandwiched between a CD63 capture antibody and enzyme coated GPC-1 and EpCAM detection antibody Figure 6A. While forming this immunocomplex, it is crucial to exercise caution during the incubation of reporter antibodies mixture to minimize the antibody crosstalk. In our case, the steps of conjugating an enzyme (beta galactosidase/ alkaline phosphate) to a detection antibody involves the incubation of biotinylated detection antibodies with streptavidin-conjugated enzymes to create a detection antibody-enzyme adduct. However, biotin and streptavidin have strong affinity for each other and can conjugate if both are incubated together. We hence incubated biotinylated detection antibodies and streptavidin conjugated enzymes sequentially during the reporter incubation step with the magnetic beads and employed three washes of magnetic beads after each incubation to minimize the cross-conjugation of enzymes and antibodies.

**Figure 6:**
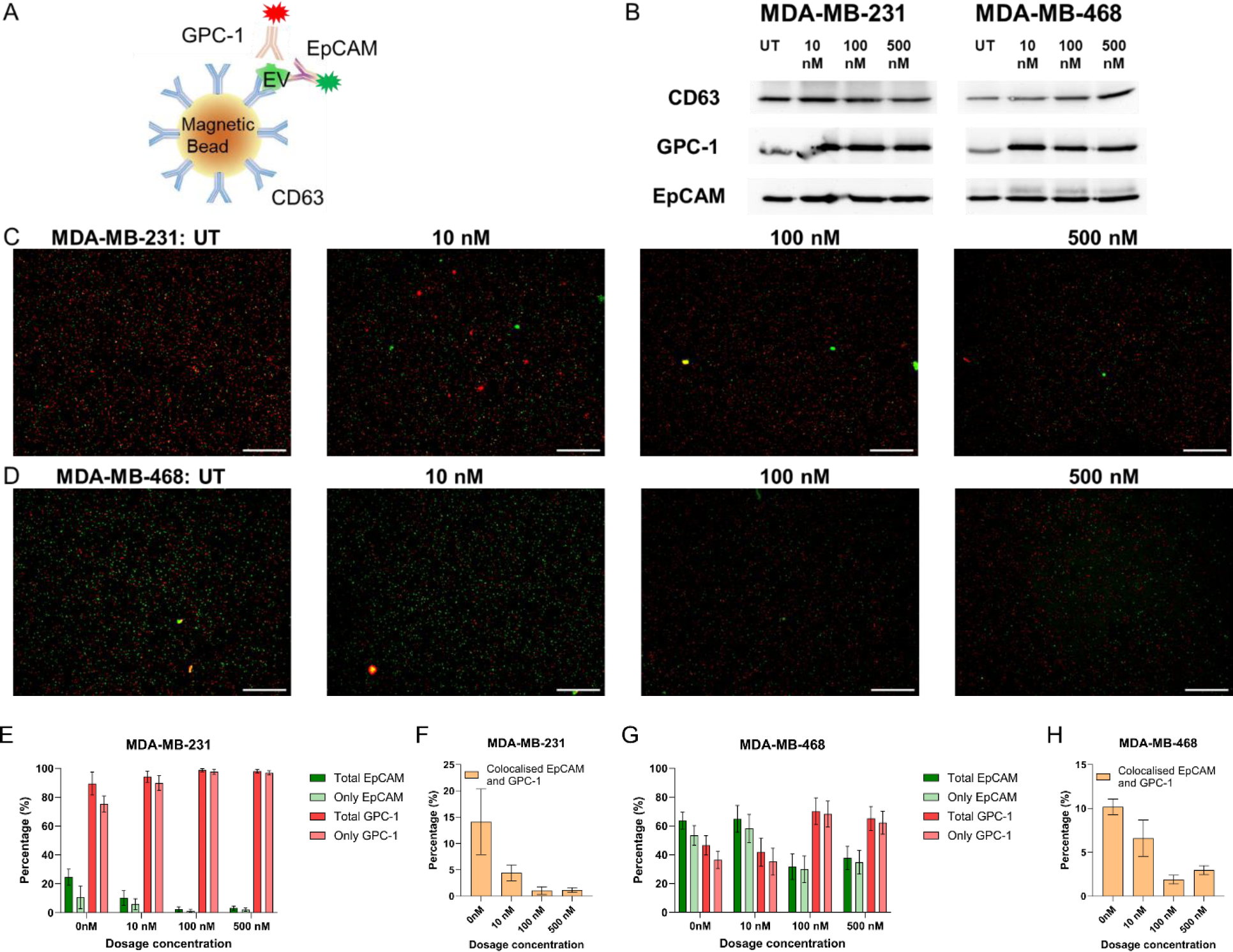
Mem-dELISA device utilization for performing single EV assay to study the effect of Paclitaxel drug treatment on breast cancer cell lines. **A)** A schematic of the immunocomplex of EV sandwiched between CD63 coated magnetic beads and two reporter antibodies (EpCAM and GPC-1) conjugated with ALP and beta galactosidase enzyme respectively. **B)** Western blot images of CD63, GPC-1 and EpCAM proteins from cell lysate as function of drug concentration (10 nM, 100 nM and 500 nM) for MDA-MB-231 and MDA-MB-468 cell lines. **C & D)** Overlayed fluorescent images (red: GPC-1 and green: EpCAM) obtained from duplex digital ELISA of EVs as a function of drug dosage for both MDA-MB-231 and MDA-MB-468 cell lines, respectively. Left to right images represent untreated control, 10 nM,100 nM and 500 nM Paclitaxel drug treatment. **E) & G)** Plot of % of EpCAM and GPC-1 protein content on EVs as function of drug concentration for MDA-MB-231 and MDA-MB-468 cell lines, respectively. **F & H)** Graph showing the % of colocalized EVs (both EpCAM and GPC-1) as a function of drug concentration for MDA-MB-231 and MDA-MB-468 cell lines, respectively. Scale bar is 200 µm.

The presence of GPC-1 and EpCAM in the cell lysate was confirmed by Western blot (Figure 6B). For characterization of the effect of Paclitaxel drug treatment on the EVs derived from breast cancer cells, first the cell lines were treated with different concentrations of paclitaxel (10 nM, 100 nM, and 500 nM). Following the drug treatment, the EVs were isolated from the cell culture media by ultracentrifugation. Figure 1C shows the complete workflow of the Mem-dELISA platform for the duplex detection of GPC-1 and EpCAM proteins after immunocomplex formation. Figure 6C and 6D show a series of representative fluorescence images from both breast cancer cell lines to elucidate the effect of drug treatment on EVs for both MDA-MB-231 and MDA-MB-468 cell lines. Images from the left to right images represent the digital assay performed on EVs which were obtained from untreated control, 10 nM, 100 nM, and 500 nM Paclitaxel drug treated cell lines. The red color corresponds to GPC-1 positive EVs while the green color corresponds to EpCAM positive EVs. The fraction of the EVs belonging to GPC-1 only, EPCAM only and colocalized (both GPC-1 and EpCAM) is calculated based on set theory as shown in Supplementary Figure 12. As shown in Figures 6F and 6H, for both the breast cancer cell lines, as the concentration of paclitaxel was increased, the colocalized fraction of GPC-1 and EpCAM monotonically decreased. In contrast, the fraction of GPC-1 increases with increase in drug treatment concentration (Figures 6E and 6G). However, in the case of EpCAM alone, the fraction decreases monotonically with an increase of paclitaxel. These results suggest that the drug treatment suppresses the expression of EpCAM but not GPC-1. It also indicates that cancerous cells still exist and may require higher doses of chemotherapy to suppress the expression GPC-1. Interestingly, this selective suppression cannot be identified by Western blot analysis of cell lysate as evident in Figure 6B. This is likely due to the presence of dispersed soluble proteins in the cell culture media that generate false positive signals. This result suggests that our mem-dELISA EV colocalization assay provides more tumor state-relevant information than Western blot analysis of cell lysate.

## 4. Conclusion

In this report, we have presented Mem-dELISA platform: a dual color, lithography-free, highly economical (< $0.1 consumable cost) digital biosensor which integrated a piece of commercial PCTE membrane into a microfluidics channel to form thousands of microwells. The facile integration of membrane into the microfluidics channels coupled with the wicking effect eliminated the need for expensive vacuum, fluidic pumps, or valves. For duplex color detection, two enzymatic amplification reactions namely beta-galactosidase and alkaline phosphate along with their selected substrates were individually optimized and subsequently performed simultaneously in the same experiment. Following the optimization of the bead loading efficiency to >80% through the introduction of a permanent magnet and mechanical shaking, the biosensor achieved a dynamic range of 1 pM-10 aM (5 logs) with 10 aM limit of detection for B-βG and B-ALP proteins which is comparable to commercial Simoa device. Finally, the Mem-dELISA platform was validated by performing digital ELISA of EVs extracted from two breast cancer cell lines after treatment with variable dosage of chemotherapy drug. The colocalized fraction of EpCAM and GPC-1 protein pair and the EpCAM itself decreased with the drug dosage suggesting that these two ratios can be used as a surrogate marker for assessing therapy outcomes.

The number of microwells in the Mem-dELISA device is solely contingent on the size of the membrane providing a facile means for scaling up to achieve ∼1 million microwells per chip. This scalability facilitates highly multiplexed operations. Additionally, leveraging the duplex color capability derived from two enzymatic amplification reactions, the developed digital enzyme-linked immunosorbent assay (ELISA) method exhibits substantial potential for massive multiplexing when integrated with barcoded magnetic beads in a single run.

Furthermore, augmenting the number of membrane pieces per chip (approximately 10) and establishing connections with the top microfluidics channel can empower the device to facilitate around 600 multiplexed assays(Song et al., 2021). With the additional capability of duplex colors from 2 enzymatic amplification reactions, the developed digital ELISA method can be massively multiplexed when coupled with bar coded magnetic beads in a single run. While the current testing has focused on EVs, the same workflow can be applied to detect and conduct colocalization assays for any multi-epitope-based antigens, such as viruses, lipoproteins, ribonucleoproteins, among others. Beyond digital ELISA, we anticipate that this technology can find application in digital immune-PCR, hybridization assays, single cell analysis and colony formation assay.

## Author contributions

V.Y., H.S. and H.C.C. conceived the project. H.S., V.Y. designed the experiments. H.S. and V.Y. designed, fabricated, tested, and optimized the Mem-dELISA device. H.S. and V.Y. developed the image analysis codes, performed data analysis and numerical simulations. A.B. performed cell culture, exosome extraction and western blot experiments. T.S. helped in experiments. H.C.C., S.S. and M.D. supervised the work. All authors contributed to writing the manuscript.

## Acknowledgments

This work has been supported by the National Institute of Health (NIH) Common Fund, through the Office of Strategic Coordination/Office of the NIH Director, 4UH3CA241684-03 (H.C.C. and S.S.), the National Cancer Institute grant K22CA258410 (M.D.), and the National Institute of General Medical Sciences grant R35GM151041 (M.D.). We thank Dr. Golnaz Asaadi Tehrani for her guidance regarding breast cancer chemotherapy studies, and Ms. R’nld Rumbach for technical assistance.

## Appendix A. Supplementary data

Supplementary data to this article can be found online at

## References

Alba-Bernal, A., Lavado-Valenzuela, R., Domínguez-Recio, M.E., Jiménez-Rodriguez, B., Queipo-Ortuño, M.I., Alba, E., Comino-Méndez, I., 2020. Challenges and achievements of liquid biopsy technologies employed in early breast cancer. eBioMedicine 62. 10.1016/j.ebiom.2020.103100

Ali, N.S., Akudugu, J.M., Howell, R.W., 2019. A preliminary study on treatment of human breast cancer xenografts with a cocktail of paclitaxel, doxorubicin, and 131I-anti-epithelial cell adhesion molecule (9C4). World Journal of Nuclear Medicine 18, 18–24.

Arab, T., Mallick, E.R., Huang, Y., Dong, L., Liao, Z., Zhao, Z., Gololobova, O., Smith, B., Haughey, N.J., Pienta, K.J., Slusher, B.S., Tarwater, P.M., Tosar, J.P., Zivkovic, A.M., Vreeland, W.N., Paulaitis, M.E., Witwer, K.W., 2021. Characterization of extracellular vesicles and synthetic nanoparticles with four orthogonal single-particle analysis platforms. Journal of Extracellular Vesicles 10, e12079. 10.1002/jev2.12079

Chang, L., Rissin, D.M., Fournier, D.R., Piech, T., Patel, P.P., Wilson, D.H., Duffy, D.C., 2012. Single Molecule Enzyme-Linked Immunosorbent Assays: Theoretical Considerations. J Immunol Methods 378, 102–115. 10.1016/j.jim.2012.02.011

Ciravolo, V., Huber, V., Ghedini, G.C., Venturelli, E., Bianchi, F., Campiglio, M., Morelli, D., Villa, A., Mina, P.D., Menard, S., Filipazzi, P., Rivoltini, L., Tagliabue, E., Pupa, S.M., 2012. Potential role of HER2-overexpressing exosomes in countering trastuzumab-based therapy. Journal of Cellular Physiology 227, 658–667. 10.1002/jcp.22773

Cohen, L., Walt, D.R., 2019. Highly Sensitive and Multiplexed Protein Measurements. Chem. Rev. 119, 293–321. 10.1021/acs.chemrev.8b00257

Connal, S., Cameron, J.M., Sala, A., Brennan, P.M., Palmer, D.S., Palmer, J.D., Perlow, H., Baker, M.J., 2023. Liquid biopsies: the future of cancer early detection. J Transl Med 21, 118. 10.1186/s12967-023-03960-8

Dutt, S., Apel, P., Lizunov, N., Notthoff, C., Wen, Q., Trautmann, C., Mota-Santiago, P., Kirby, N., Kluth, P., 2021. Shape of nanopores in track-etched polycarbonate membranes. Journal of Membrane Science 638, 119681. 10.1016/j.memsci.2021.119681

Hinestrosa, J.P., Kurzrock, R., Lewis, J.M., Schork, N.J., Schroeder, G., Kamat, A.M., Lowy, A.M., Eskander, R.N., Perrera, O., Searson, D., Rastegar, K., Hughes, J.R., Ortiz, V., Clark, I., Balcer, H.I., Arakelyan, L., Turner, R., Billings, P.R., Adler, M.J., Lippman, S.M., Krishnan, R., 2022. Early-stage multi-cancer detection using an extracellular vesicle protein-based blood test. Commun Med 2, 1–9. 10.1038/s43856-022-00088-6

Kelley, S.O., Mirkin, C.A., Walt, D.R., Ismagilov, R.F., Toner, M., Sargent, E.H., 2014. Advancing the speed, sensitivity and accuracy of biomolecular detection using multi-length-scale engineering. Nature nanotechnology 9, 969–980.

Kelly, K.M., Dean, J., Lee, S.-J., Comulada, W.S., 2010. Breast cancer detection: radiologists’ performance using mammography with and without automated whole-breast ultrasound. Eur Radiol 20, 2557–2564. 10.1007/s00330-010-1844-1

Landegren, U., Hammond, M., 2021. Cancer diagnostics based on plasma protein biomarkers: hard times but great expectations. Molecular Oncology 15, 1715–1726. 10.1002/1878-0261.12809

Li, W., Shao, B., Liu, C., Wang, H., Zheng, W., Kong, W., Liu, X., Xu, G., Wang, C., Li, H., Zhu, L., Yang, Y., 2018. Noninvasive Diagnosis and Molecular Phenotyping of Breast Cancer through Microbead-Assisted Flow Cytometry Detection of Tumor-Derived Extracellular Vesicles. Small Methods 2, 1800122. 10.1002/smtd.201800122

Lim, C.T., Zhang, Y., 2007. Bead-based microfluidic immunoassays: The next generation. Biosensors and Bioelectronics 22, 1197–1204. 10.1016/j.bios.2006.06.005

Lin, X., Huang, X., Urmann, K., Xie, X., Hoffmann, M.R., 2019. Digital loop-mediated isothermal amplification on a commercial membrane. ACS sensors 4, 242–249.

Liu, C., Xu, X., Li, B., Situ, B., Pan, W., Hu, Y., An, T., Yao, S., Zheng, L., 2018. Single-Exosome-Counting Immunoassays for Cancer Diagnostics. Nano Lett. 18, 4226–4232. 10.1021/acs.nanolett.8b01184

Liu, X., Li, J., Cadilha, B.L., Markota, A., Voigt, C., Huang, Z., Lin, P.P., Wang, D.D., Dai, J., Kranz, G., Krandick, A., Libl, D., Zitzelsberger, H., Zagorski, I., Braselmann, H., Pan, M., Zhu, S., Huang, Y., Niedermeyer, S., Reichel, C.A., Uhl, B., Briukhovetska, D., Suárez, J., Kobold, S., Gires, O., Wang, H., 2019. Epithelial-type systemic breast carcinoma cells with a restricted mesenchymal transition are a major source of metastasis. Science Advances 5, eaav4275. 10.1126/sciadv.aav4275

Lu, N., Zhang, M., Lu, L., Liu, Y.-Z., Zhang, H.-H., Liu, X.-D., 2021. miRNA-490-3p promotes the metastatic progression of invasive ductal carcinoma. Oncology Reports 45, 706–716. 10.3892/or.2020.7880

Matsuda, K., Maruyama, H., Guo, F., Kleeff, J., Itakura, J., Matsumoto, Y., Lander, A.D., Korc, M., 2001. Glypican-1 Is Overexpressed in Human Breast Cancer and Modulates the Mitogenic Effects of Multiple Heparin-binding Growth Factors in Breast Cancer Cells1. Cancer Research 61, 5562–5569.

Morasso, C., Ricciardi, A., Sproviero, D., Truffi, M., Albasini, S., Piccotti, F., Sottotetti, F., Mollica, L., Cereda, C., Sorrentino, L., Corsi, F., 2022. Fast quantification of extracellular vesicles levels in early breast cancer patients by Single Molecule Detection Array (SiMoA). Breast Cancer Res Treat 192, 65–74. 10.1007/s10549-021-06474-3

Obayashi, Y., Iino, R., Noji, H., 2015. A single-molecule digital enzyme assay using alkaline phosphatase with a cumarin-based fluorogenic substrate. Analyst 140, 5065–5073. 10.1039/C5AN00714C

Ono, T., Ichiki, T., Noji, H., 2018. Digital enzyme assay using attoliter droplet array. Analyst 143, 4923–4929. 10.1039/C8AN01152D

Rissin, D.M., Kan, C.W., Campbell, T.G., Howes, S.C., Fournier, D.R., Song, L., Piech, T., Patel, P.P., Chang, L., Rivnak, A.J., Ferrell, E.P., Randall, J.D., Provuncher, G.K., Walt, D.R., Duffy, D.C., 2010. Single-molecule enzyme-linked immunosorbent assay detects serum proteins at subfemtomolar concentrations. Nat Biotechnol 28, 595–599. 10.1038/nbt.1641

Rissin, D.M., Kan, C.W., Song, L., Rivnak, A.J., Fishburn, M.W., Shao, Q., Piech, T., Ferrell, E.P., Meyer, R.E., Campbell, T.G., Fournier, D.R., Duffy, D.C., 2013. Multiplexed single molecule immunoassays. Lab Chip 13, 2902. 10.1039/c3lc50416f

Rivnak, A.J., Rissin, D.M., Kan, C.W., Song, L., Fishburn, M.W., Piech, T., Campbell, T.G., DuPont, D.R., Gardel, M., Sullivan, S., 2015. A fully-automated, six-plex single molecule immunoassay for measuring cytokines in blood. Journal of immunological methods 424, 20–27.

Roganovic, D., Djilas, D., Vujnovic, S., Pavic, D., Stojanov, D., 2015. Breast MRI, digital mammography and breast tomosynthesis: Comparison of three methods for early detection of breast cancer. Bosn J Basic Med Sci 15, 64–68. 10.17305/bjbms.2015.616

Rondelez, Y., Tresset, G., Tabata, K.V., Arata, H., Fujita, H., Takeuchi, S., Noji, H., 2005. Microfabricated arrays of femtoliter chambers allow single molecule enzymology. Nat Biotechnol 23, 361–365. 10.1038/nbt1072

Sakakihara, S., Araki, S., Iino, R., Noji, H., 2010. A single-molecule enzymatic assay in a directly accessible femtoliter droplet array. Lab Chip 10, 3355. 10.1039/c0lc00062k

Schwarzenbach, H., Gahan, P.B., 2020. Predictive value of exosomes and their cargo in drug response/resistance of breast cancer patients. Cancer Drug Resist 3, 63–82. 10.20517/cdr.2019.90

Shao, H., Im, H., Castro, C.M., Breakefield, X., Weissleder, R., Lee, H., 2018. New Technologies for Analysis of Extracellular Vesicles. Chem. Rev. 118, 1917–1950. 10.1021/acs.chemrev.7b00534

Sharma, H., John, K., Gaddam, A., Navalkar, A., Maji, S.K., Agrawal, A., 2018. A magnet-actuated biomimetic device for isolating biological entities in microwells. Sci Rep 8, 12717. 10.1038/s41598-018-31274-z

Sharma, H., Yadav, V., D’Souza-Schorey, C., Go, D.B., Senapati, S., Chang, H.-C., 2023. A Scalable High-Throughput Isoelectric Fractionation Platform for Extracellular Nanocarriers: Comprehensive and Bias-Free Isolation of Ribonucleoproteins from Plasma, Urine, and Saliva. ACS Nano 17, 9388–9404. 10.1021/acsnano.3c01340

Shim, J., Ranasinghe, R.T., Smith, C.A., Ibrahim, S.M., Hollfelder, F., Huck, W.T.S., Klenerman, D., Abell, C., 2013. Ultrarapid Generation of Femtoliter Microfluidic Droplets for Single-Molecule-Counting Immunoassays. ACS Nano 7, 5955–5964. 10.1021/nn401661d

Song, Y., Ye, Y., Su, S.-H., Stephens, A., Cai, T., Chung, M.-T., K. Han, M., W. Newstead, M., Yessayan, L., Frame, D., David Humes, H., H. Singer, B., Kurabayashi, K., 2021. A digital protein microarray for COVID-19 cytokine storm monitoring. Lab on a Chip 21, 331–343. 10.1039/D0LC00678E

Soysal, S.D., Muenst, S., Barbie, T., Fleming, T., Gao, F., Spizzo, G., Oertli, D., Viehl, C.T., Obermann, E.C., Gillanders, W.E., 2013. EpCAM expression varies significantly and is differentially associated with prognosis in the luminal B HER2+, basal-like, and HER2 intrinsic subtypes of breast cancer. Br J Cancer 108, 1480–1487. 10.1038/bjc.2013.80

Tian, F., Zhang, S., Liu, C., Han, Z., Liu, Yuan, Deng, J., Li, Y., Wu, X., Cai, L., Qin, L., Chen, Q., Yuan, Y., Liu, Yi, Cong, Y., Ding, B., Jiang, Z., Sun, J., 2021. Protein analysis of extracellular vesicles to monitor and predict therapeutic response in metastatic breast cancer. Nat Commun 12, 2536. 10.1038/s41467-021-22913-7

Verbruggen, B., Tóth, T., Cornaglia, M., Puers, R., Gijs, M.A.M., Lammertyn, J., 2015. Separation of magnetic microparticles in segmented flow using asymmetric splitting regimes. Microfluid Nanofluid 18, 91–102. 10.1007/s10404-014-1409-8

Wei, P., Wu, F., Kang, B., Sun, X., Heskia, F., Pachot, A., Liang, J., Li, D., 2020. Plasma extracellular vesicles detected by Single Molecule array technology as a liquid biopsy for colorectal cancer. Journal of Extracellular Vesicles 9, 1809765. 10.1080/20013078.2020.1809765

Wills, C.A., Liu, X., Chen, L., Zhao, Y., Dower, C.M., Sundstrom, J., Wang, H.-G., 2021. Chemotherapy-Induced Upregulation of Small Extracellular Vesicle-Associated PTX3 Accelerates Breast Cancer Metastasis. Cancer Research 81, 452–463. 10.1158/0008-5472.CAN-20-1976

Wilson, D.H., Rissin, D.M., Kan, C.W., Fournier, D.R., Piech, T., Campbell, T.G., Meyer, R.E., Fishburn, M.W., Cabrera, C., Patel, P.P., Frew, E., Chen, Y., Chang, L., Ferrell, E.P., von Einem, V., McGuigan, W., Reinhardt, M., Sayer, H., Vielsack, C., Duffy, D.C., 2016. The Simoa HD-1 Analyzer: A Novel Fully Automated Digital Immunoassay Analyzer with Single-Molecule Sensitivity and Multiplexing. J Lab Autom. 21, 533–547. 10.1177/2211068215589580

Witters, D., Knez, K., Ceyssens, F., Puers, R., Lammertyn, J., 2013. Digital microfluidics-enabled single-molecule detection by printing and sealing single magnetic beads in femtoliter droplets. Lab on a Chip 13, 2047–2054. 10.1039/C3LC50119A

Wu, C., Dougan, T.J., Walt, D.R., 2022. High-Throughput, High-Multiplex Digital Protein Detection with Attomolar Sensitivity. ACS Nano 16, 1025–1035. 10.1021/acsnano.1c08675

Wu, C., Maley, A.M., Walt, D.R., 2020. Single-molecule measurements in microwells for clinical applications. Critical Reviews in Clinical Laboratory Sciences 57, 270–290. 10.1080/10408363.2019.1700903

Yadav, V., Chong, N., Ellis, B., Ren, X., Senapati, S., Chang, H.-C., Zorlutuna, P., 2020. Constant-potential environment for activating and synchronizing cardiomyocyte colonies with on-chip ion-depleting perm-selective membranes. Lab Chip 20, 4273–4284. 10.1039/D0LC00809E

Yu, S., Wei, Y., Xu, Y., Zhang, Y., Li, J., Zhang, J., 2016. Extracellular vesicles in breast cancer drug resistance and their clinical application. Tumor Biol. 37, 2849–2861. 10.1007/s13277-015-4683-5

Zandi Shafagh, R., Decrop, D., Ven, K., Vanderbeke, A., Hanusa, R., Breukers, J., Pardon, G., Haraldsson, T., Lammertyn, J., van der Wijngaart, W., 2019. Reaction injection molding of hydrophilic-in-hydrophobic femtolitre-well arrays. Microsyst Nanoeng 5, 25. 10.1038/s41378-019-0065-2

