## Supplementary Files for "A Mem-dELISA platform for dual color and ultrasensitive digital detection of colocalized proteins on extracellular vesicles"

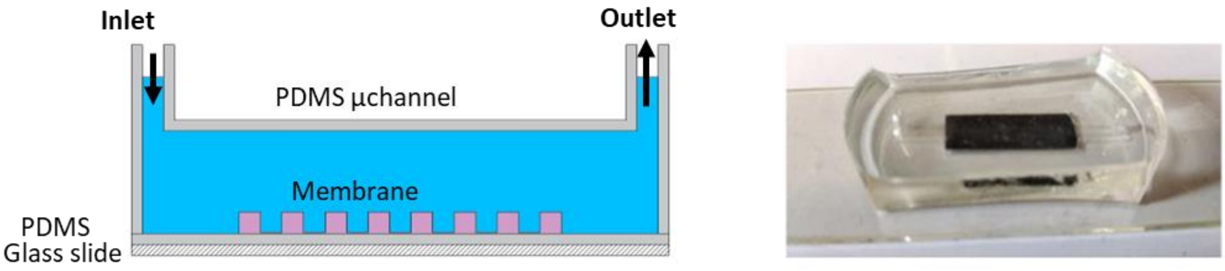


**Supplementary Figure 1:** Schematics and image of the final assembled Mem-dELISA device.

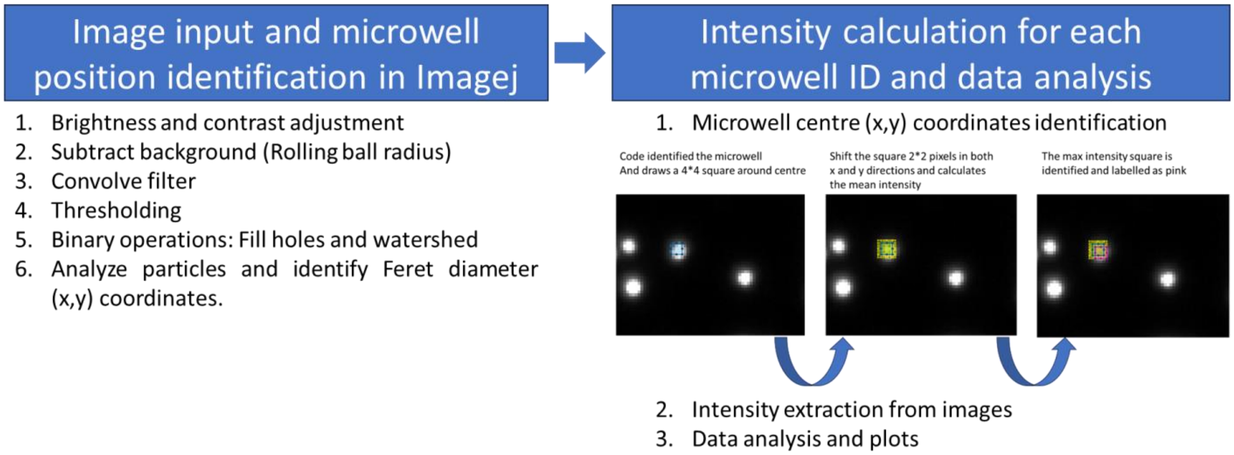


**Supplementary Figure 2:** The image processing involves an initial step aimed at noise reduction and the preliminary determination of the approximate coordinates of the microwell center. Following the identification of these coordinates, a 4x4 pixel square, depicted in blue, is generated as illustrated in the leftmost image. This square undergoes a translational shift of up to 2 pixels in both the X and Y directions to obtain the maximum microwell intensity, thereby facilitating accurate determination of the microwell coordinates. The pink square, characterized by the maximum intensity, is subsequently selected as the focal region for acquiring the precise (x, y) coordinates of the microwell center.

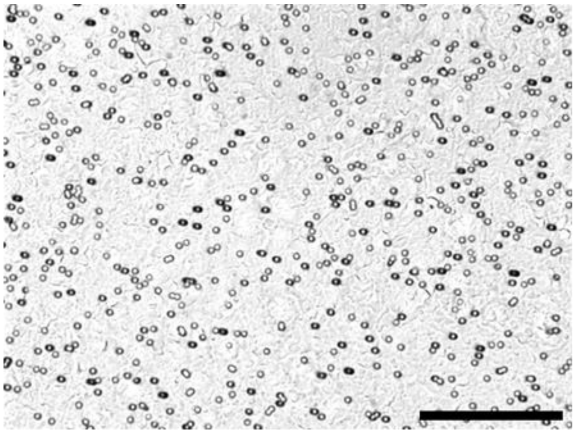

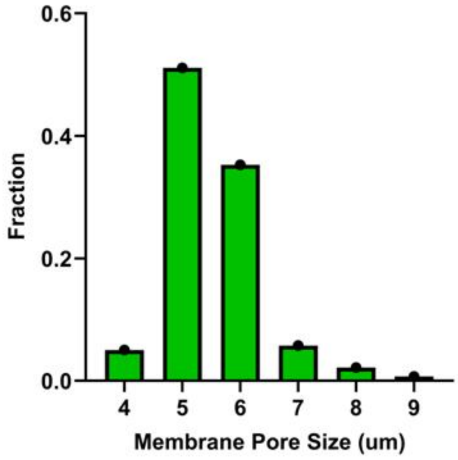


**Supplementary Figure 3**: Bright field image of the PCTE membrane and the distribution of membrane pore sizes.

| Enzyme 1 and Substrate | Enzyme 2 and Substrate | Results |
|---|---|---|
| Horseradish Peroxidase (HRP) with QuantaRed™ enhanced chemifluorescent HRP substrate | Beta-galactosidase with FDG substrate | Enzyme cross talk with substrates observed. |
| ALP with ImmPACT® Vector® Red Substrate Kit | Beta-galactosidase with FDG substrate | Enzyme cross talk with substrates observed. Beta galactosidase reaction was inhibited. |
| ALP with AttoPhos AP Fluorescent Substrate | Beta-galactosidase with FDG substrate | Enzyme cross talk with substrates observed. Beta galactosidase reaction was inhibited. |
| ALP with 4MUP substrate | Beta-galactosidase with FDG substrate | Fluorescence overlap |
| ALP with 4MUP substrate | Beta-galactosidase with RDG substrate | No enzyme crosstalk observed and this enzyme substrates combination was selected for duplex digital protein detection. |

**Supplementary Figure 4**: Analysis of various enzyme-substrate combinations to ascertain the combination exhibiting minimal fluorescence crosstalk and inhibitory behavior.

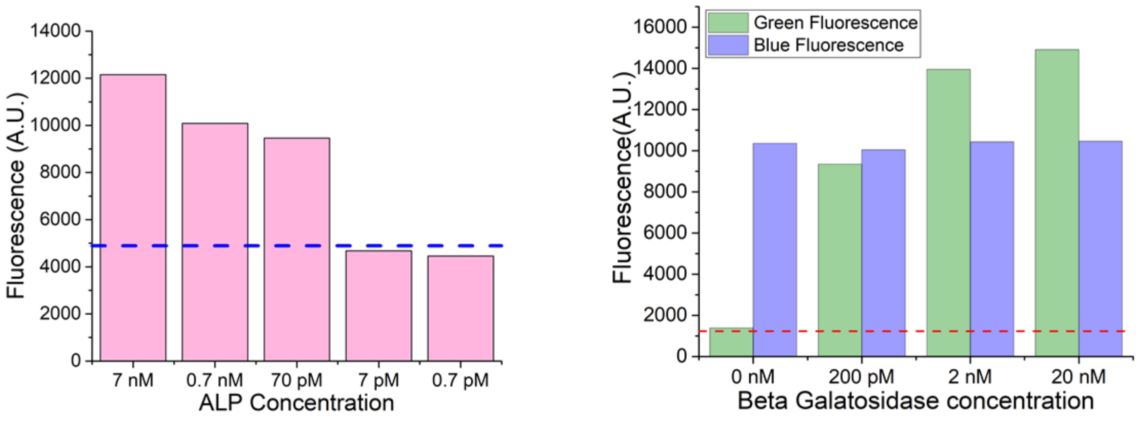


**Supplementary Figure 5:** *Left*: Bulk results of ALP amplification as a function of concentration using 2x diluted 4-MUP substrate. *Right*: Bulk amplification of beta galactosidase at various concentration while keeping the ALP concentration fixed at 70 pM. For this reaction 4-MUP and RDG substrates were mixed together.

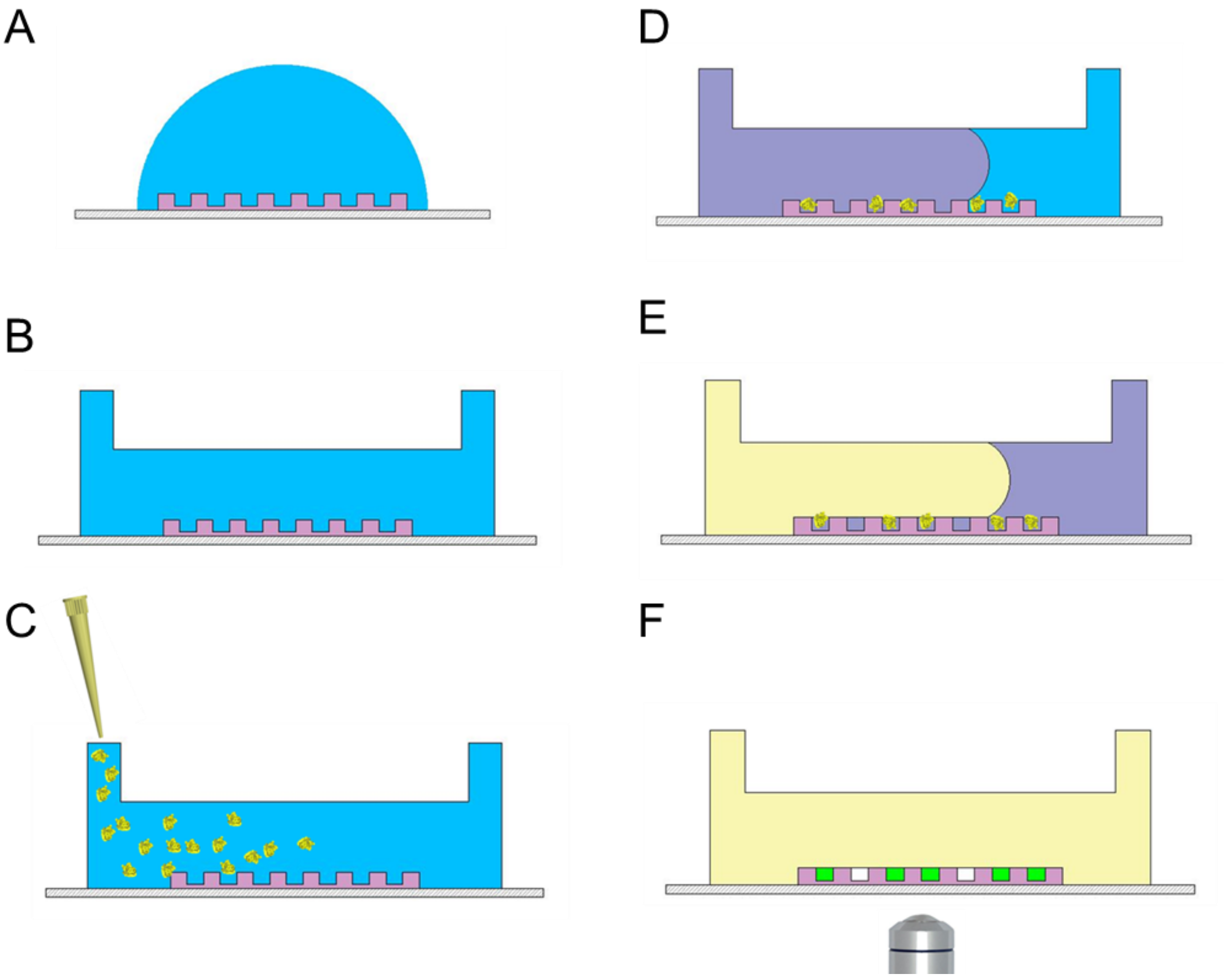


**Supplementary Figure 6: (A-F)** Workflow of random encapsulation of B-βG or B-ALP proteins followed by substrates injection and oil sealing

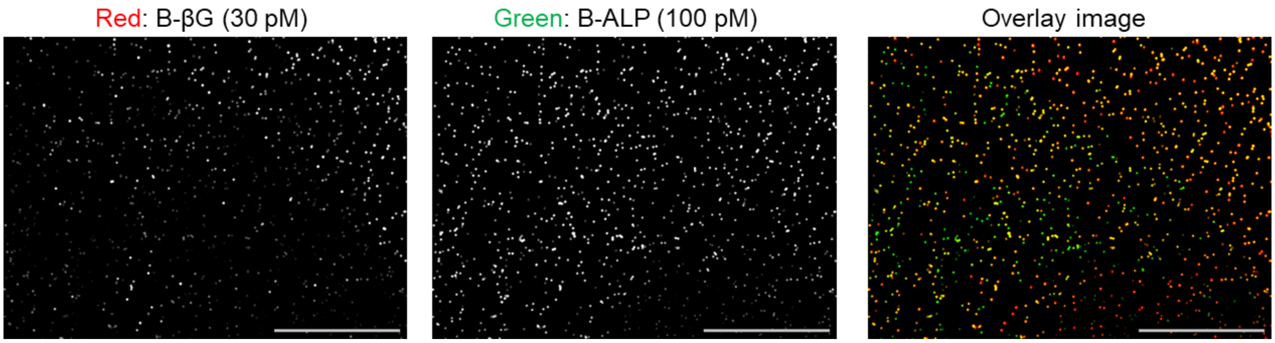


**Supplementary Figure 7:** Random encapsulation of B-βG or B-ALP molecules inside the microwells using 30 pM B-βG and 100 pM B-ALP molecules.

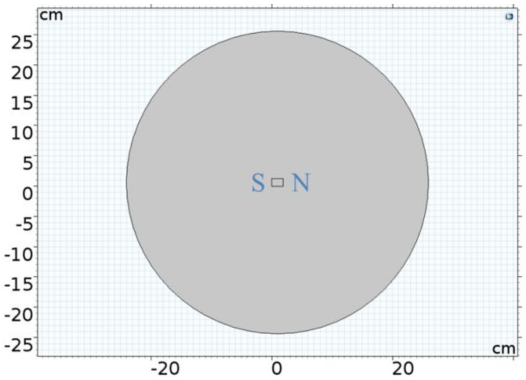


**Supplementary Figure 8**: Domain geometry for simulating a permanent magnet with magnetization of 750 kA/m.

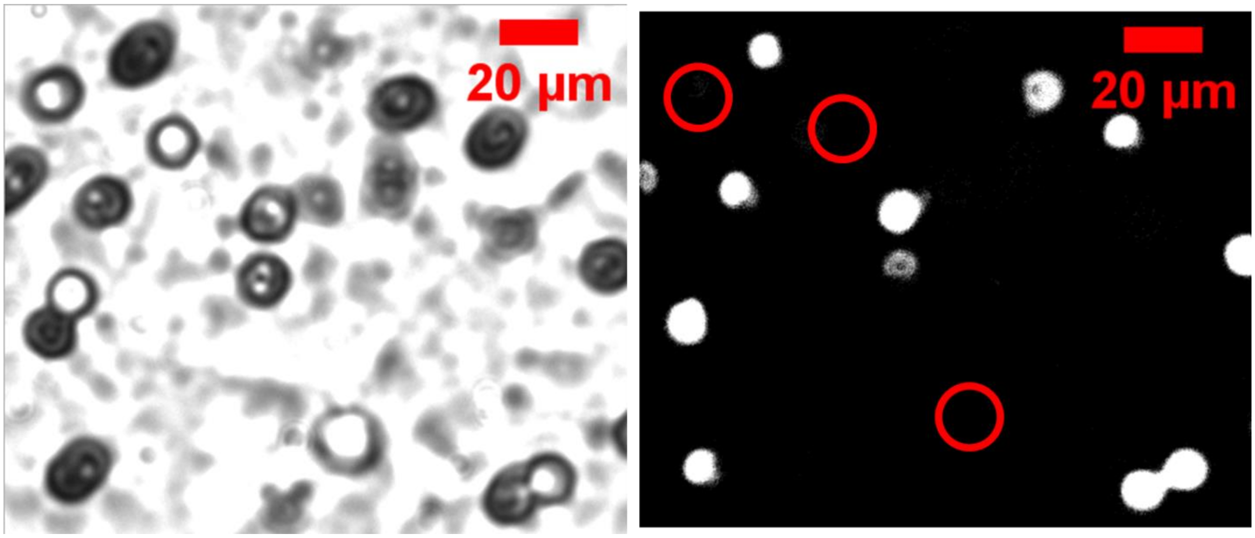


**Supplementary Figure 9:** Bright field and its corresponding fluorescence image at 50x resolution confirming that only the magnetic bead containing microwells were in 'on' state

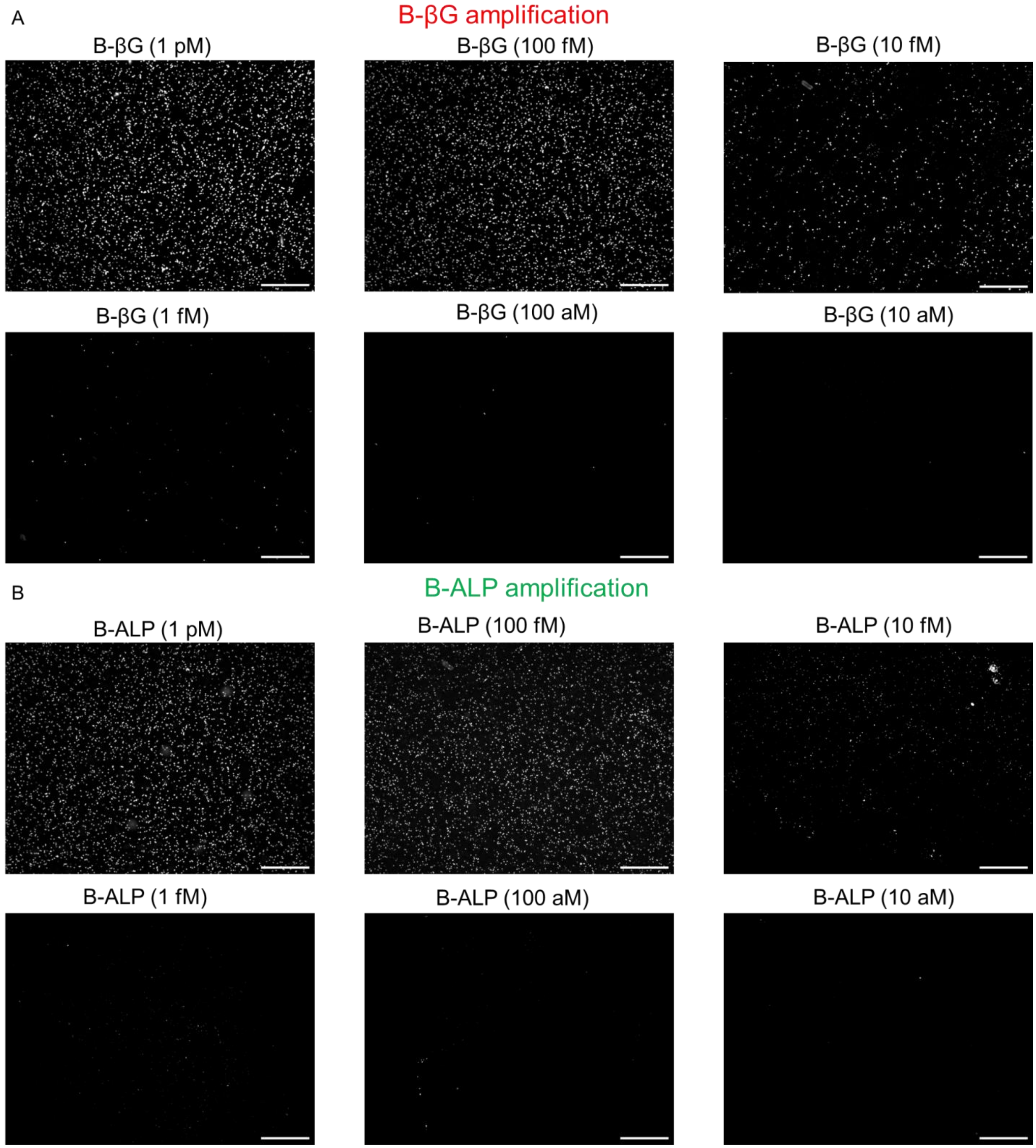


**Supplementary Figure 10:** Representative images of digital assay with serial dilution in concentration (1 pM – 10 aM) for (A) B-βG and (B) B-ALP.

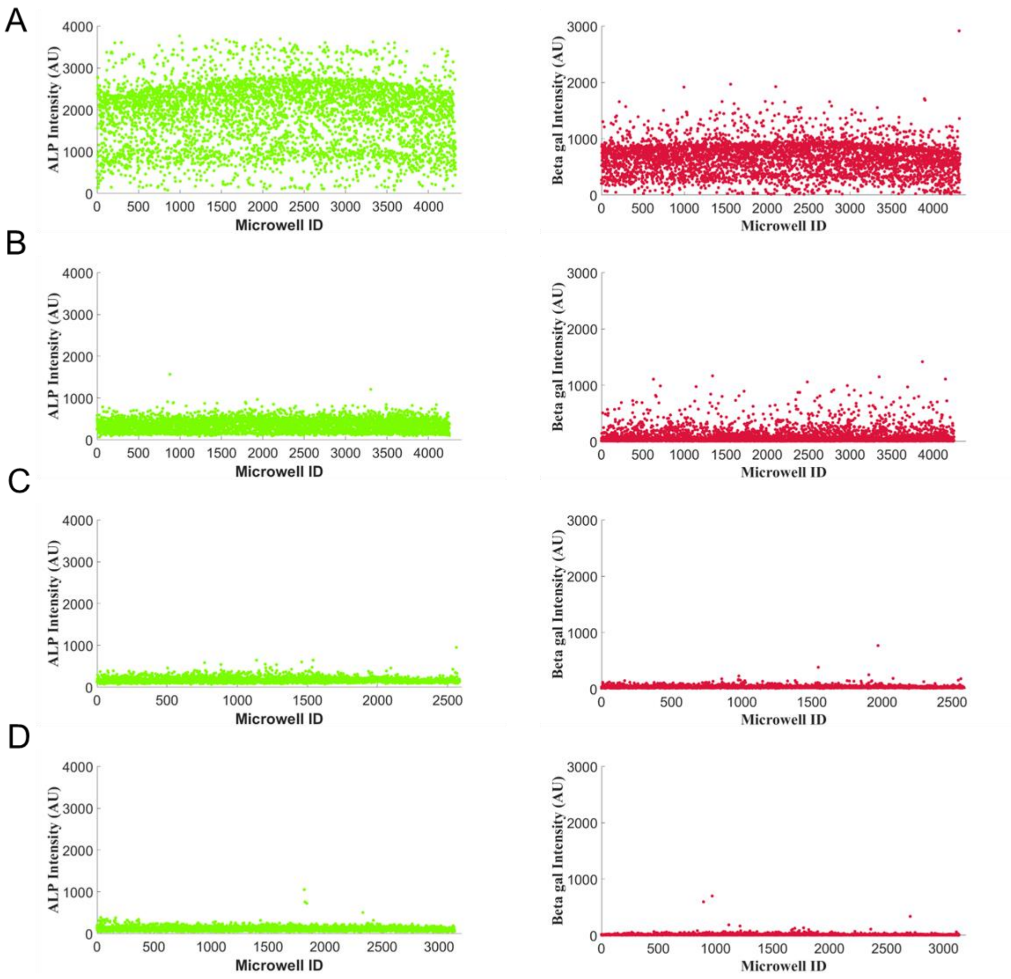


**Supplementary Figure 11:** Representative scatter plots of fluorescence intensity (B-ALP (green) and B-βG (red)) of the microwell vs microwell ID for digital assay. The sample concentration was serially diluted while keeping the B-ALP to B-βG ratio constant at 10. The B-ALP & B-βG pair concentration varied from (A) 1 pM-100 fM, (B) 100fM-10fM, (C) 10 fM-1 fM and (D) 1fM-100 aM respectively.

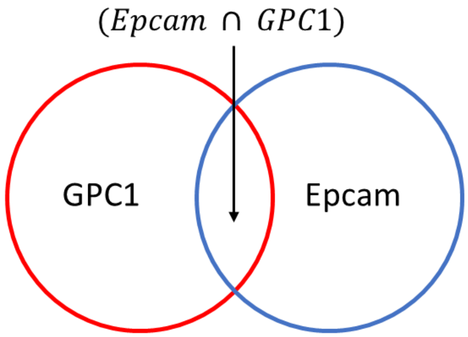


$$Total\ Epcam = \frac{Epcam}{(Epcam \cup GPC1)} \qquad Only\ Epcam = \frac{Epcam - (Epcam \cap GPC1)}{(Epcam \cup GPC1)}$$

$$Total\ GPC1 = \frac{GPC1}{(Epcam \cup GPC1)} \qquad Only\ GPC1 = \frac{GPC1 - (Epcam \cap GPC1)}{(Epcam \cup GPC1)}$$

$$Both\ Epcam\ and\ GPC1 = \frac{(Epcam \cap GPC1)}{(Epcam \cup GPC1)}$$

**Supplementary Figure 12:** Normalization scheme for protein co-localization assay for GPC-1 and Epcam proteins present on EVs using set theory.

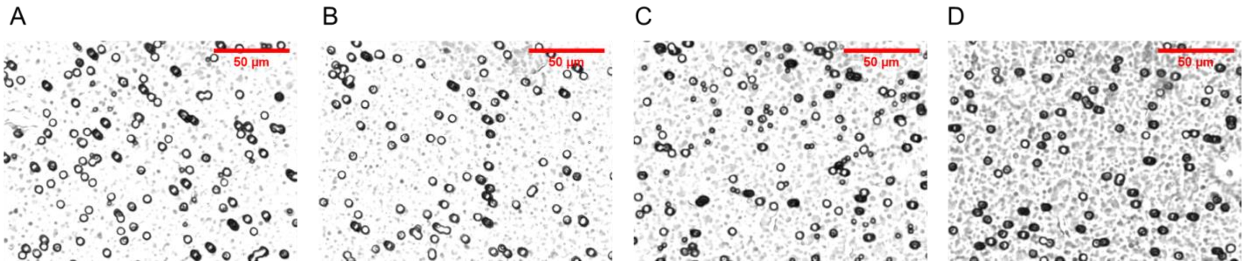


**Supplementary Figure 13:** Images depicting the filling of microwells by using (A) $2 * 10^7 beads/ml$ (B) $2.4 * 10^7 beads/ml$ and (C) $1 * 10^8 beads/ml$. (D) Image showing increase in bead filling at $2.4 * 10^7 beads/ml$ concentration using an external permanent magnet coupled with mechanical shaking.
